# Naïve Bayes taxonomic classification cost in QIIME 2 is set by the invocation rather than by sequencing depth or community richness at single-sample scale: a measured resource protocol for 16S rRNA pipelines

**DOI:** 10.64898/2026.09.01.748377

**Authors:** Sam Victor, Siddharth Sadasivam, Gugan Satheesh, Shravan Aarya Prabhakaran, S Yuvaraj, R Subha

**Affiliations:** Sri SriEshwar College of Engineering, Coimbatore, Tamil Nadu, India

**Author notes:** Correspondence: Sam Victor.

**Keywords:** 16S rRNA, amplicon sequencing, QIIME 2, DADA2, taxonomic classification, parallel scaling

## Abstract

Marker-gene amplicon workflows run on shared computing infrastructure, where cores, memory and wall time must be requested before a job starts. Those requests are set by convention, and the convention assumes a pipeline scales with the size of its data. For one of the two dominant stages, it does not.

We measured a QIIME 2 16S rRNA workflow across nine upper-respiratory samples from a paediatric otitis media cohort. Over a 27-fold range of sequencing depth, DADA2 denoising rose 12.8-fold in wall time while Naïve Bayes classification changed by one per cent across samples of low to moderate richness, 5 to 70 ASVs each and 218 across the pooled cohort. Richness did not saturate, so the invariance is not saturation: classification is dominated by a fixed cost paid before any sequence is examined. Profiling that cost shows it is compute-bound rather than storage-bound, and nearly half of it is the framework importing plugins the job never calls: classifying in an environment holding only the plugins the step needs is 43.6% faster, uses 28% less memory and returns a byte-identical result. Parallelism returns little for denoising and makes classification slower, though running samples concurrently raises throughput 21-fold against one job running alone.

Sequences and taxonomy are unchanged across thread counts, so the implied savings cost nothing scientifically. The magnitude of the fixed cost is a property of the installation and the reference artifact rather than a constant of the method, so the protocol, measurements and code are released for the constants to be remeasured where they are to be relied on.

## 1 INTRODUCTION

Amplicon sequencing of marker genes is the standard approach for culture-free profiling of microbial communities, and QIIME 2 (Bolyen et al., 2019) with DADA2 (Callahan et al., 2016) and a Naïve Bayes classifier trained against SILVA (Quast et al., 2013; Bokulich et al., 2018) forms the computational core of many 16S rRNA workflows, increasingly run on shared resources managed by schedulers such as SLURM (Yoo et al., 2003). How much resource to request is left to the user and is generally decided by heuristic: requests that are too small cause jobs to be killed mid-run, requests that are too large idle cores and lengthen queues. Workflow managers (Di Tommaso et al., 2017; Köster and Rahmann, 2012) and the amplicon pipelines built on them (Ewels et al., 2020; Straub et al., 2020; Thompson et al., 2022; Saenz et al., 2022; Cousson et al., 2025) solve task orchestration but do not predict how long a task will take or how much memory it will need.

The cost of choosing badly is measurable. Classification of a single sample is fastest at one job: requesting forty returns the same result 1.28 times slower while consuming 10.5 times the CPU. Denoising rewards threads only weakly, two already delivering 93% of the best speedup observed at any count. The saving is free to take: the reduced allocation returns representative sequences and taxonomic assignments identical at 1, 4 and 40 threads.

Runtime for these stages is not unmeasured. Bokulich et al. (2018) fit runtime as a linear function of query and reference sequence counts, recovering 23 ms per query over sets of up to 10 000 sequences, and Thompson et al. (2022) report wall times for a complete workflow on 96 samples. Both measure where per-sequence work is substantial; a single sample presents tens of representative sequences, and what dominates the cost in that range is what we address.

There is a structural reason to expect the usual mental model to be wrong. A 16S pipeline is commonly described as scaling with “the size of the data”, implicitly the number of reads, but the two expensive stages do not receive the same input: denoising consumes the full read set, while classification consumes only the amplicon sequence variants (ASVs) it returns, together with a reference whose size is fixed and independent of the study. An ASV is a complete distinct amplicon sequence, not a variant position within one: two reads differing at a single nucleotide anywhere along the amplicon are resolved as two ASVs, each emitted once as a representative sequence of full amplicon length, here a V4 region of about 253 bp. The ASV count of a sample is therefore the number of distinct sequences the classifier must label, and nothing else about the sample enters that stage. It is wide-ranging and community-dependent: the middle-ear effusion measured here returns five, the adenoid samples around seventy, and soil or sediment communities routinely hundreds to thousands (Thompson et al., 2017). The range spanned here, 5 to 70 ASVs per sample and 218 across the pooled cohort, sits at the low end of what the stage can be asked to do (Section 6.1).

Richness saturation with depth is well established (Gotelli and Colwell, 2001; Willis, 2019) and is the obvious mechanism to expect here. What we measure is stronger and has a different cause: richness did not saturate, rising 2.8-fold, and classification cost still did not move, because the stage is dominated by a cost the query count does not enter at all at this scale. Either way, sizing a classification job by read count measures a quantity that stage does not depend on, which is what every resource heuristic we are aware of does, including the SLURM templates previously distributed with our own pipeline.

We test this on a cohort in which richness varies for biological reasons rather than by construction: a paediatric otitis media study profiling middle ear, nasopharynx and adenoid in parallel. We use that gradient as a source of realistic computational inputs and nothing more; nine samples from six participants cannot characterise the ecology of those sites, and no claim of that kind is made.

Specifically we:

1. **Measure what each stage costs.** Denoising and classification across a 26.9-fold range of sequencing depth, between samples and within one library subsampled to six depths, separating classification into fixed and per-ASV components.
2. **Characterise how each stage scales,** from 1 to 40 physical cores, fitting Amdahl’s Law (Amdahl, 1967) and the Universal Scalability Law (Gunther, 2007).
3. **Decompose the fixed cost and remove part of it.** Nearly half is the framework importing plugins the job never calls: classifying in a minimal environment is 43.6% faster and uses 28% less memory while returning a byte-identical result.
4. **Verify that the result is unchanged,** so that a reduced allocation is a saving rather than a trade.
5. **Turn the measurements into recommendations**, and release the instrumentation, the measured data and the analysis code.

What follows is written as a protocol: the constants are specific to this system, the relationships are not.

## 2 MATERIALS AND EQUIPMENT

### 2.1 Computing environment

All measurements were made on a single dual-socket workstation: two Intel Xeon Gold 6230 processors (2 sockets *×* 20 cores = 40 physical cores, 80 logical), 251 GiB RAM, running Ubuntu 22.04.5 (kernel 5.15.0). Local storage was a single ext4 volume. No other substantial workload ran concurrently.

The protocol itself requires only a multi-core Linux machine with QIIME 2 installed. The reported constants are hardware-specific; the procedure is not, and Section 3.5 states which quantities are which.

### 2.2 Sequence data

All sequence data are previously published and publicly archived. We used nine runs from European Nucleotide Archive study PRJEB33591 (ERP116397), an investigation of the middle-ear microbiome in paediatric otitis media with effusion (Jörissen et al., 2021). Sampling one study throughout removes batch, extraction-protocol and primer-variant differences as confounders, which matters because ASV count is the dependent variable here and is sensitive to all three.

All runs are Illumina MiSeq, paired-end 2 *×* 251 bp, generated with the Caporaso 515F/806R primer pair (Caporaso et al., 2011) and deposited already primer-trimmed; revised forms of this pair exist (Parada et al., 2016; Apprill et al., 2015; Walters et al., 2016) and differ in degeneracy, so we state the variant explicitly. Measured median merged ASV length was 253 bp throughout. Table 1 lists the runs, their depths and the resulting richness.

**Table 1.** Sequence data. All runs from PRJEB33591; Illumina MiSeq, 2 *×* 251 bp, 16S rRNA V4 (515F/806R). ASVs are the count after DADA2 denoising at `--p-trunc-len-f 240 --p-trunc-len-r 240`. The nine runs come from six participants: D-003, D-006 and D-008 each contribute two samples from different sites, so the runs are not fully independent, which is why the cost-against-richness comparison of Section 4.4 is reported as an effect size and a correlation described as descriptive rather than as a significance test. The ASV counts in the final column are per sample, before pooling; the deduplicated union across the nine samples is 218 distinct ASVs, which is the query set used wherever the pooled cohort is measured.

| **Run** | **Participant** | **Source** | **Read pairs** | **ASVs** |
|---|---|---|---|---|
| ERR3444623 | D-005 | middle ear | 62 444 | 5 |
| ERR3444633 | D-006 | middle ear | 54 938 | 11 |
| ERR3444597 | D-002 | nasopharynx | 70 630 | 16 |
| ERR3444606 | D-003 | ear canal | 44 604 | 38 |
| ERR3444605 | D-003 | nasopharynx | 66 993 | 54 |
| ERR3444642 | D-008 | nasopharynx | 66 212 | 55 |
| ERR3444628 | D-006 | nasopharynx | 1 804 054 | 58 |
| ERR3444680 | D-014 | adenoid | 62 686 | 67 |
| ERR3444641 | D-008 | adenoid | 82 257 | 70 |

The depth comparison uses two nasopharyngeal samples, ERR3444605 (66 993 read pairs) and ERR3444628 (1 804 054 read pairs), from the same study, protocol and body site but *different participants* (D-003 and D-006), so composition varies alongside depth and this contrast alone cannot separate the two. We therefore also subsample a single library across a range of depths (Section 3.7).

Both reads were truncated at 240 bp: a symmetric cut at that length retains an expected overlap of about 227 bp between mates, far above the 12 bp DADA2 requires to merge, while removing the final low-quality bases. Holding one value across all nine libraries keeps ASV count comparable between them, which is the quantity the study depends on; this is a cost study rather than an ecological one, so the truncation was chosen to be defensible and uniform rather than tuned per library.

### 2.3 Pipeline and software

The pipeline is a six-stage Bash driver: retrieval from the ENA, import into QIIME 2, DADA2 denoising, taxonomic classification, artifact export and visualisation. Only denoising and classification are computationally significant. Measurements were made under QIIME 2 2026.7 (`qiime2`, `rachis` and `q2cli` all 2026.7.0) with `q2-dada2` 2026.7.0 wrapping `bioconductor-dada2` 1.38.0, scikit-learn 1.7.1 and Python 3.12, on Ubuntu 22.04.5 (kernel 5.15.0).

The contents of that environment are a parameter of the results and not background detail, because Section 4.5 shows that nearly half the fixed classification cost is the framework importing the plugins it finds installed. The `rachis` distribution for this release specifies 683 packages, among them 27 QIIME 2 plugins and 235 R or Bioconductor packages reached through `rpy2`, which `q2-dada2` requires and classification does not use. A reader comparing classification timings against ours should compare installed plugin sets as carefully as they compare hardware. A full `conda` export is distributed with the code, but as a provenance record rather than an installation recipe: every dependency carries a platform-specific build hash, so readers reproducing this work should install the official `rachis` distribution for the corresponding release, or better a lockfile or container image. For the minimal classification environment of Section 4.6 this repository now ships all three: `environments/qiime2-lean.yml`, an installation recipe carrying version pins but no build hashes; `environments/Dockerfile.lean`; and `environments/qiime2-lean.def` for Apptainer at sites without Docker.

The distinction is not academic. The pipeline as originally published targeted QIIME 2 2025.7, and four interface changes required code modification before it would run at all under 2026.7: manifest-referenced FASTQ files must now be gzipped, `dada2 denoise-paired` requires an additional `--o-base-transition-stats` output, `feature-classifier extract-reads` requires `--o-read-extraction-stats`, and the distribution itself was renamed from `amplicon` to `rachis`, so the documented installation command no longer resolves. A fifth is reported in Section 2.4. Stating the version alone is not sufficient to make a result reproducible.

### 2.4 Reference classifier

The classifier was trained locally rather than downloaded: QIIME 2 no longer distributes region-specific classifiers, the last SILVA 515F/806R artifact it hosted was serialised under scikit-learn 0.24.1 and is refused by current releases, and from 2026.4 the project refers users to external providers whose available builds were documented against earlier release ranges. We trained it with RESCRIPt (Robeson et al., 2021) against SILVA 138.2 SSU NR99 (Quast et al., 2013; Yilmaz et al., 2014), extracting V4 with the same Caporaso 515F (GTGCCAGCMGCCGCGGTAA) and 806R (GGACTACHVGGGTWTCTAAT) primers used to generate the data, dereplicating in `uniq` mode, and fitting a multinomial Naïve Bayes model (Wang et al., 2007; Bokulich et al., 2018) — the configuration practitioners actually run, and so the one whose cost is worth measuring. The training script is released and is deterministic given the SILVA release and the primer pair, so a reader can rebuild the artifact rather than transfer it.

QIIME 2 also ships alignment-based classifiers (`classify-consensus-vsearch`, `-blast`), which we did not benchmark and would not expect the central result to carry over to: their cost is dominated by alignment work that scales with the number of queries, where `classify-sklearn` pays most of its cost before the first query is examined. The fixed-cost finding is specific to the Naïve Bayes path. No claim about assignment quality is made anywhere in this work; accuracy comparisons are a separate and well-covered question (Bokulich et al., 2018; Straub et al., 2020; Prodan et al., 2020).

The artifact is 62 MiB on disk, reported because a reader comparing timings should know it. Section 4.5 quantifies how much the total depends on it: 2.2 s to read and deserialise it and a further 4.50 s in a scoring call whose cost is set by the 674 MiB likelihood matrix rather than by the queries, so roughly 6.6 s of the stage — a fifth of the cost in the full distribution and about a third once the unused plugins are removed (Section 4.6). Artifact size and installed plugin set are both confounds when comparing installations, the second being the larger.

### 2.5 Instrumentation and analysis code

The measurement harness, the pipeline it drives, the resource predictor and the figure-generation scripts are distributed together (Data Availability Statement). Five components carry the protocol. `lib/benchmark.sh` wraps a stage in `/usr/bin/time -v` and appends one row per run to a CSV. `scripts/run real benchmarks.sh` implements the experimental arms of Section 3.7. `scripts/prepare datasets.sh` stages and denoises the input runs, and `scripts/build classifier.sh` trains the reference classifier. `lib/predictor.py` fits the scaling and memory models.

Each stage is invoked under `/usr/bin/time -v` with the linear-algebra thread pools pinned (`OMP NUM THREADS`, `OPENBLAS NUM THREADS`, `MKL NUM THREADS`, `NUMEXPR NUM THREADS` and `VECLIB MAXIMUM THREADS` all set to 1), so that the stage’s own thread parameter is the only source of parallelism. The two measured commands are

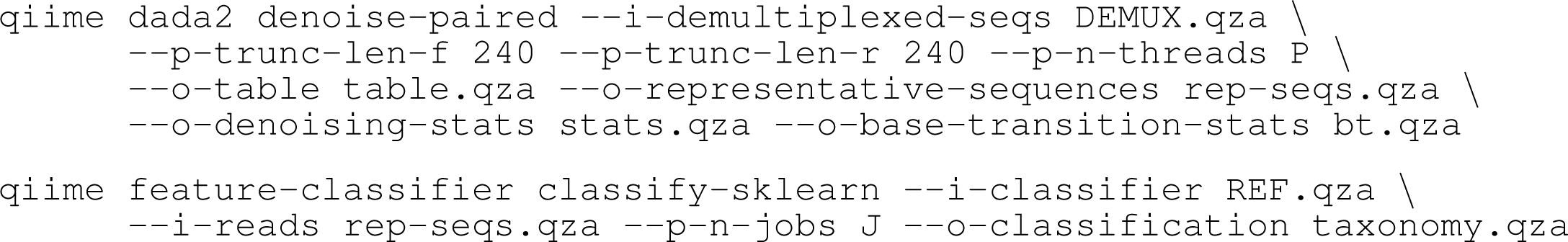

with `P` and `J` varied across the ladder of Section 3.7. Any run whose recorded exit status is non-zero is discarded before fitting, for the reason given in Section 3.4.

## 3 METHODS

### 3.1 Objective

The protocol measures wall-clock time and peak memory for the two dominant stages of a QIIME 2 amplicon pipeline and separates the two input dimensions they respond to: read count for denoising, ASV count for classification. Its output is a set of stage-specific resource models for sizing scheduler requests, together with the measurements they were fitted to.

### 3.2 Protocol

Table 2 gives the seven steps with the command that performs each, their approximate duration on the hardware of Section 2.1, and the two points at which the procedure must be halted and an intermediate result checked. Every step is also a runnable script in the project repository, https://github.com/samvictordr/16S-Amplicon-rRNA-pipeline-protocol. Both pause points guard against failures that produce no error at all: a classifier trained on the wrong primer region still produces taxonomy, and a dataset whose reads retain their PCR primers still produces ASVs.

**Table 2.**
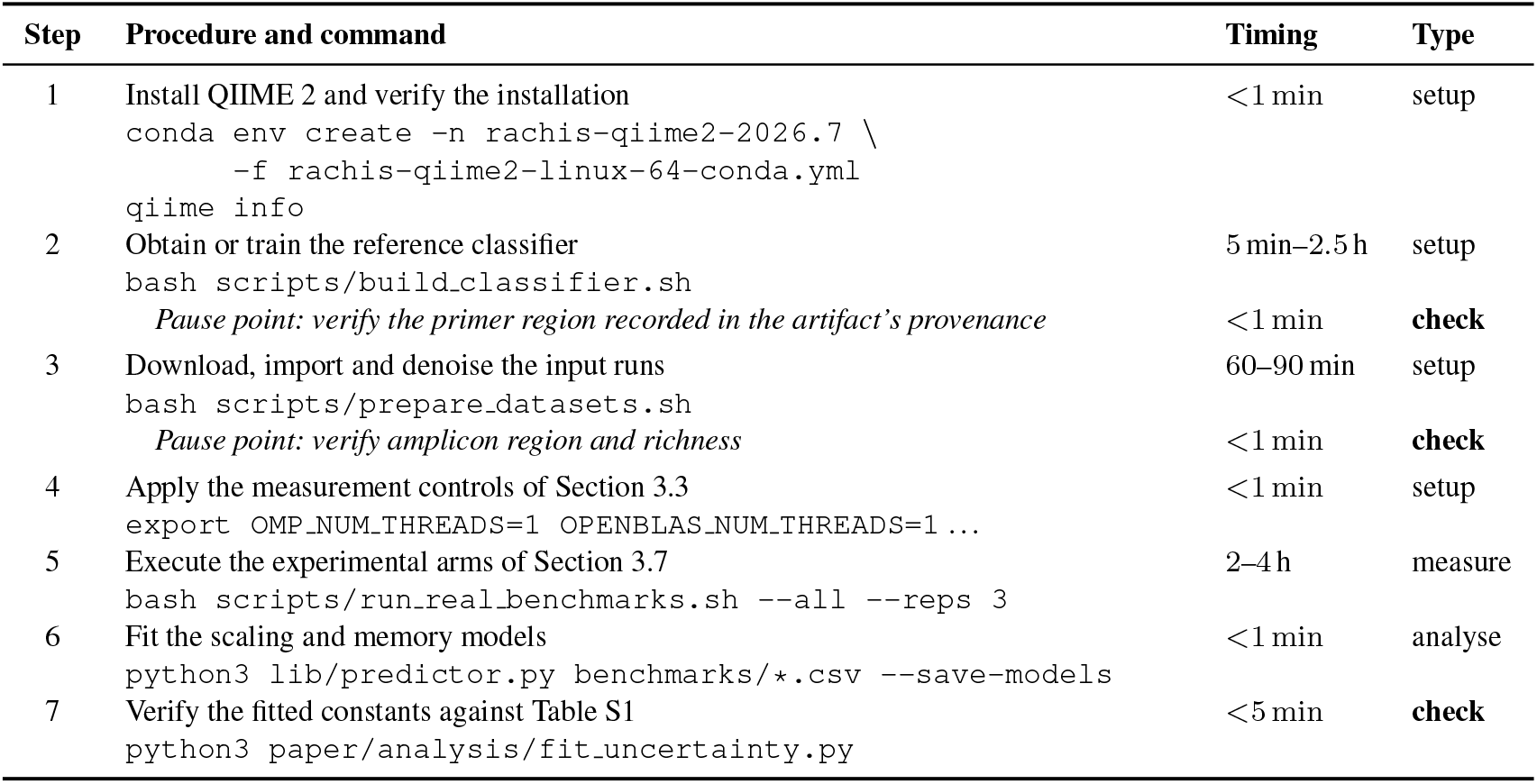
Protocol steps, with the command that performs each. Durations are for the hardware in Section 2.1 with 40 cores available and are dominated by download bandwidth in steps 2–3. Pause points are checkpoints at which an intermediate result must be verified before continuing; expected values for every measured quantity are given in Table S1.

#### Step 1: Environment

Install the QIIME 2 distribution for the release in use, named rachis from 2026.7, and confirm the version; the rename from amplicon is one of the interface changes Table S2 warns about.

#### Step 2: Reference Classifier

Obtain a Naïve Bayes classifier whose extracted region matches the amplicon, downloading a compatible pre-trained artifact where one exists and otherwise training one (Section 2.4). *Pause point.* Read the primer pair back out of the artifact’s provenance and confirm it is either the amplicon’s own region or an untrimmed full-length reference; a full-length classifier is a legitimate choice, a classifier extracted for a *different* region is not. This check exists because an earlier revision of this pipeline classified V4 reads against a V3–V4 classifier for an extended period without any error being raised.

#### Step 3: Input Data

Download, import and denoise each input run, recording read count and resulting ASV count. *Pause point.* Confirm that the median merged ASV length is consistent with the target amplicon (*∼*253 bp for V4) before continuing. A median near 292 bp indicates reads that retain their PCR primers (253 + 19 + 20); because primer positions are degenerate, DADA2 resolves the primer variants as biological variation and inflates the ASV count, which is the dependent variable here.

#### Steps 4–7

Apply the controls of Section 3.3 — this takes no appreciable time and is the step most easily omitted — then execute the experimental arms of Section 3.7 at three repetitions each, or ten where small differences are at stake, fit the models of Section 3.8 to the resulting CSVs, and check the fitted constants and key ratios against Section 3.5.

### 3.3 Measurement controls and their validation

Each stage invocation was wrapped in `/usr/bin/time -v`, recording wall time, CPU time, peak resident set size (in kibibytes, so all memory figures here are binary multiples), context switches and filesystem I/O counts. Most points were repeated three times and we report the mean; the decomposition, the randomised replication and the plugin-set comparison used ten and the cold-cache arms five, as Table S3 states. The sampling profile of Section 4.5 was collected with `py-spy` 0.4.2 at 50 Hz; the profiler slows the invocation it samples by roughly 6*×*, so it is used to attribute time proportionally and never to measure it.

Two terms are not interchangeable: a *thread* is what DADA2 spawns through `--p-n-threads`, sharing one address space, where a *job* is what `classify-sklearn` spawns through `--p-n-jobs`, separate `loky` worker processes that must each be started and must return their results. Two controls were applied. NumPy and SciPy spawn BLAS thread pools independently of the thread count requested from QIIME 2; these were pinned to one thread, without which a nominally single-threaded denoising run consumed 105 s of CPU in 58 s of wall time, understating every subsequent speedup. Second, core counts were enumerated from physical cores via `lscpu`.

Machine-level settings were left at distribution defaults, and two bear on the results. No NUMA policy was applied, so the scheduler could migrate processes between the two nodes of 125.6 GiB each; we now regard that as the one control we should have applied, since placing a classification’s CPU on one node and its memory on the other costs 4.82 s, 14.4% of the stage, where binding both to the same node costs nothing measurable. Anyone repeating this on a multi-socket machine should bind both with `numactl --cpunodebind=N --membind=N` and report that they did. Second, turbo boost was enabled against a 2.10 GHz base clock, so a run at one core and a run at 40 are not at the same clock and part of the strong-scaling decline is frequency rather than coordination overhead. We have not isolated that part, but wall and CPU time diverge by an order of magnitude across the ladder, which no clock change can produce; it bears on the magnitude but not the sign, and not at all on the depth and richness results, measured at a fixed four cores. Runs were made with a warm page cache, the cold penalty being measured separately at 1.90 s, about 5%.

### 3.4 Advantages, pitfalls and troubleshooting

Table S2 lists the pitfalls encountered during this work with the symptom that reveals each and its remedy. The first is the largest: left unpinned, a nominally single-threaded denoising run consumed 105 s of CPU in 58 s of wall time, and every speedup computed against such a baseline is understated.

### 3.5 Verification

A reader repeating this protocol on other hardware should not expect the absolute times reported here: wall times, the fitted *T*_1_, the fixed classification cost and the memory intercept are properties of the hardware, the filesystem, the installed plugin set and the reference artifact. What should transfer are the dimensionless quantities — denoising wall time rising by roughly an order of magnitude across a 27-fold depth contrast while classification moves by a few per cent (here 12.8*×* against 1.01*×*); classification speedup below 1.0 at every job count above one; classification cost near unity across an order of magnitude of richness; and bit-identical output across thread counts — together with the fraction attributable to fixed cost, which should scale chiefly with the installed plugin set and processor speed and secondarily with the reference. That is an expectation from one node and one reference, not a demonstrated result. Expected values for every measured quantity, with the file each derives from, are in Table S1. Table 3 states what each reported figure for the fixed cost is, so the small differences between the six independent determinations — which span 36.05 s to 36.49 s, a spread of 1.2% comparable to the stage’s own run-to-run scatter — are not mistaken for inconsistency.

**Table 3.**
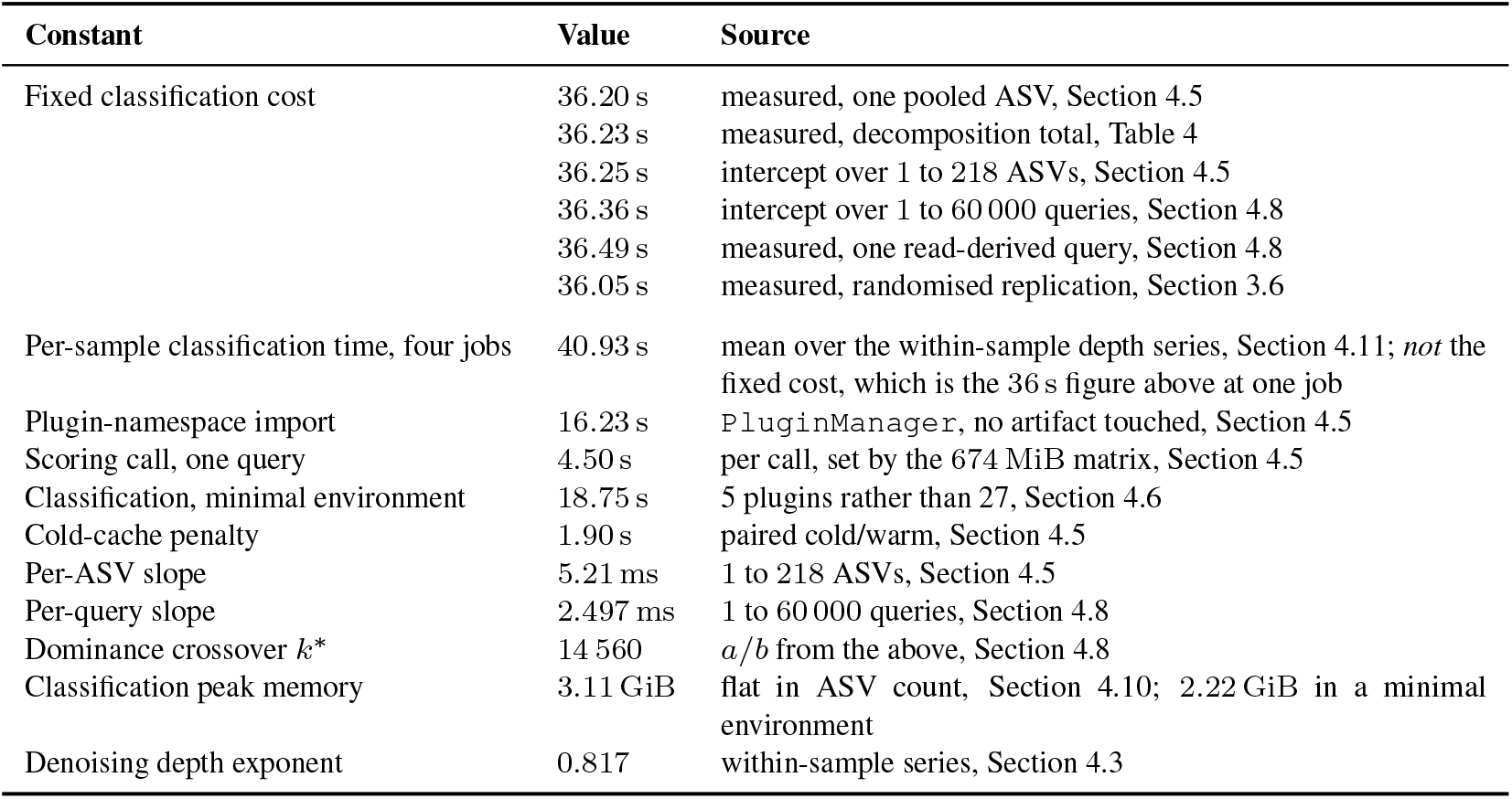
Key constants, what each one is, and where it comes from. The first six rows are independent determinations of the same quantity and agree to within 1.2%.

| Constant | Value | Source |
|---|---|---|
| Fixed classification cost | 36.20 s | measured, one pooled ASV, Section 4.5 |
|  | 36.23 s | measured, decomposition total, Table 4 |
|  | 36.25 s | intercept over 1 to 218 ASVs, Section 4.5 |
|  | 36.36 s | intercept over 1 to 60 000 queries, Section 4.8 |
|  | 36.49 s | measured, one read-derived query, Section 4.8 |
|  | 36.05 s | measured, randomised replication, Section 3.6 |
| Per-sample classification time, four jobs | 40.93 s | mean over the within-sample depth series, Section 4.11; *not* the fixed cost, which is the 36 s figure above at one job |
| Plugin-namespace import | 16.23 s | PluginManager, no artifact touched, Section 4.5 |
| Scoring call, one query | 4.50 s | per call, set by the 674 MiB matrix, Section 4.5 |
| Classification, minimal environment | 18.75 s | 5 plugins rather than 27, Section 4.6 |
| Cold-cache penalty | 1.90 s | paired cold/warm, Section 4.5 |
| Per-ASV slope | 5.21 ms | 1 to 218 ASVs, Section 4.5 |
| Per-query slope | 2.497 ms | 1 to 60 000 queries, Section 4.8 |
| Dominance crossover *k*<sup>\*</sup> | 14 560 | *a*/*b* from the above, Section 4.8 |
| Classification peak memory | 3.11 GiB | flat in ASV count, Section 4.10; 2.22 GiB in a minimal environment |
| Denoising depth exponent | 0.817 | within-sample series, Section 4.3 |

### 3.6 Replication under randomised ordering

The measurements above were taken arm by arm in a fixed order, which leaves open the possibility that an apparent effect of thread count is instead drift in machine state. We re-measured the strong scaling and ASV sweep grids eight weeks later in fully randomised order, ten repetitions per cell, 280 measurements shuffled into a single sequence rather than blocked by condition. All completed without error.

Agreement with the original triplets is close: across the eight denoising cells the largest shift in mean wall time is 0.21 s (0.43%) and seven of eight bootstrap intervals span zero; across the twelve sweep cells the largest is 0.43 s (1.04%), seven of twelve spanning zero, and the five that exclude zero all differ by less than 1.1%. The more informative quantity is the scatter: had fixed ordering been introducing drift, randomising would redistribute it within cells and inflate the within-cell spread. It did not, mean within-cell standard deviation moving from 0.189 s to 0.216 s for denoising and from 0.224 s to 0.233 s for the sweep.

The replication’s classification arm used the full 218-sequence pool where the original used 54, so those cells are not like-for-like. Normalised to each run’s own single-threaded time the thread-count response diverges by at most 0.053*×* at 40 cores and classification is slowest at the highest core count in both. The offset is itself predictable: the replicated sweep puts one sequence at 36.05 s (SD 0.11 s) — the sixth independent determination of the fixed cost in Table 3 — and 218 at 37.31 s, an implied 5.78 ms per sequence over its own endpoints, which predicts 37.63 s against 37.43 s measured, an error of 0.53%.

### 3.7 Experimental design

Eighteen experimental arms were run, listed with their repetition counts and held-fixed quantities in Table S3. Arms that vary input size hold allocation fixed at four cores and scaling arms hold input fixed; the two classifier sweeps each vary input size at both one and eight jobs, since the question they address is whether the two settings differ as a function of input size. The ASV sweep pools representative sequences from all nine samples and deduplicates exactly, giving 218 distinct sequences from 374. The extended query sweep is a computational extension and not a biological one: its queries are dereplicated reads carrying sequencing error, drawn at an even stride through the pool because reads come off the flowcell in tile order, and the taxonomy they receive is neither a result nor recorded. It overlaps the ASV sweep at 1, 10, 50, 100 and 218 sequences.

### 3.8 Scaling models

For strong scaling we fit Amdahl’s Law (Amdahl, 1967),

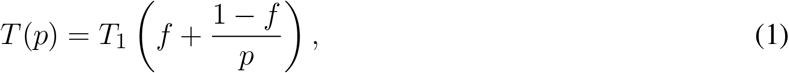

with serial fraction *f*, and the Universal Scalability Law (Gunther, 2007),

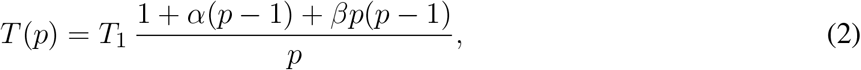

where *α* is a contention coefficient and *β* a coherency coefficient. Equation 1 is monotonic in *p* and so cannot represent a workload that becomes slower with additional cores; Equation 2 can, and admits an optimum at 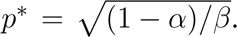 Both were fitted by non-linear least squares and compared by *R*^2^, by adjusted *R*^2^ and by AIC*_c_* (Akaike, 1974; Hurvich and Tsai, 1989), the latter two penalising the extra degree of freedom the Universal Scalability Law carries.

### 3.9 Precision and resolvable effect size

Repeatability differs markedly between the stages. Denoising’s run-to-run standard deviation is 0.05 s to 0.36 s on means of 47 s to 57 s, a coefficient of variation below 0.8%; classification’s reaches 1.15 s, 2.7%. Since every null comparison here concerns classification, that is the figure which governs what can be resolved. Repetitions were run consecutively within a batch, which understates uncertainty across batches: classification of ERR3444628 at four cores appears in three arms whose means span 0.54 s, the appropriate error term for between-arm comparisons.

The probe session of Section 4.5 bounds that term more sharply, and less comfortably. The single-sequence classification the replication had measured at 36.05 s returned 32.98 s to 33.47 s across eleven runs in three later sessions on the same idle machine, in an otherwise identical invocation: about 7% faster, against a within-cell standard deviation of 0.11 s. The between-session term is more than an order of magnitude larger than the within-session term, and it governs how far any single number here should be trusted.

We tested six candidate causes and report them in full in Supplementary Note S1. Five are excluded: the timing harness, processor frequency state, eviction of the reference by intervening work, a software change between sessions, and — the most natural explanation, and wrong — page-cache state of the import path, which reproduces the magnitude but not the persistence, recovering completely after a single run where the replication held its value for ten. The survivor is NUMA placement: no binding policy was applied, and forcing a cross-node placement costs 4.82 s (38.28 s against 33.46 s unbound) where binding both to one node costs nothing. It is larger than the 2.75 s to be explained, persists for a session because cached library pages are placed once when first read, and falls on classification rather than denoising. We cannot establish retrospectively where the replication’s pages sat and do not assert this was the cause; the effect is real, several times the within-session scatter, and binding is the control we would apply first and did not (Section 3.3).

Two consequences follow. The agreement of the six determinations of the fixed cost to 1.2% (Table 3) holds within the sessions that produced them and is not 1.2% accuracy for a value measured afresh; and a reader should expect classification timings to move by several per cent between sittings on one machine.

Three repetitions therefore resolve effects of roughly 4% and larger. This figure is a property of the instrument, fixed before any null comparison was made — the run-to-run coefficient of variation propagated to a difference of means at *n* = 3 — and not chosen with reference to any effect it is later used to judge. The ten repetitions of Section 3.6 tighten it to about 0.5% for the two arms they cover, which is why the replication rather than the original triplets is the basis for statements about small differences. It is adequate for the effects the protocol is designed to detect — the depth response of denoising being a factor of 12.8, the degradation of classification under parallelism 28% — and inadequate for establishing exact equality, so the null comparisons are reported accordingly. The practical floor is the fixed per-invocation cost itself, 36.2 s here.

## 4 RESULTS

Unless stated otherwise, timings below are means of three repetitions with BLAS thread pools pinned and core counts taken from physical cores, as described in Section 3.3. Several arms depart from this: the fixed-cost decomposition (Section 4.5), the randomised replication (Section 3.6) and the plugin-set comparison (Section 4.6) use ten repetitions, and the cold-cache arms use five paired runs.

### 4.1 Community structure across the cohort

The nine samples span a clear gradient in richness and in Shannon evenness (Shannon, 1948), computed on the per-sample feature table after denoising (Figure S1, Table S4). Middle-ear effusions are least diverse (5–38 ASVs, *H^′^* = 0.01–1.13), dominated by *Alloiococcus* or *Haemophilus*; nasopharyngeal swabs intermediate (16–58 ASVs, *H^′^* = 1.01–1.49) and dominated by *Moraxella* throughout; adenoid samples most diverse (67–70 ASVs, *H^′^* = 2.58–2.62) with *Haemophilus* at much lower dominance. One effusion (ERR3444623) is effectively clonal: 5 ASVs, 99.95% *Haemophilus*, *H^′^* = 0.005. This gradient is the axis along which classification cost is claimed not to vary, measured in real communities rather than constructed by subsetting.

All composition reported here is at genus level, which is as deep as this design resolves: of the 54 representative sequences from ERR3444605, 52 received a genus assignment and none a species name, which is expected for a 253 bp V4 amplicon where distinct species frequently share an identical fragment. Mean assignment confidence was 0.969. We make no species-level claim anywhere in this work.

### 4.2 Classification cost shows no detectable response to sequencing depth

The two nasopharyngeal samples differ 26.9-fold in sequencing depth while their ASV richness differs by 7%, which separates the two candidate input sizes cleanly. Table S5 gives the measurements and Figure 1 shows them for both stages.

**Figure 1.**
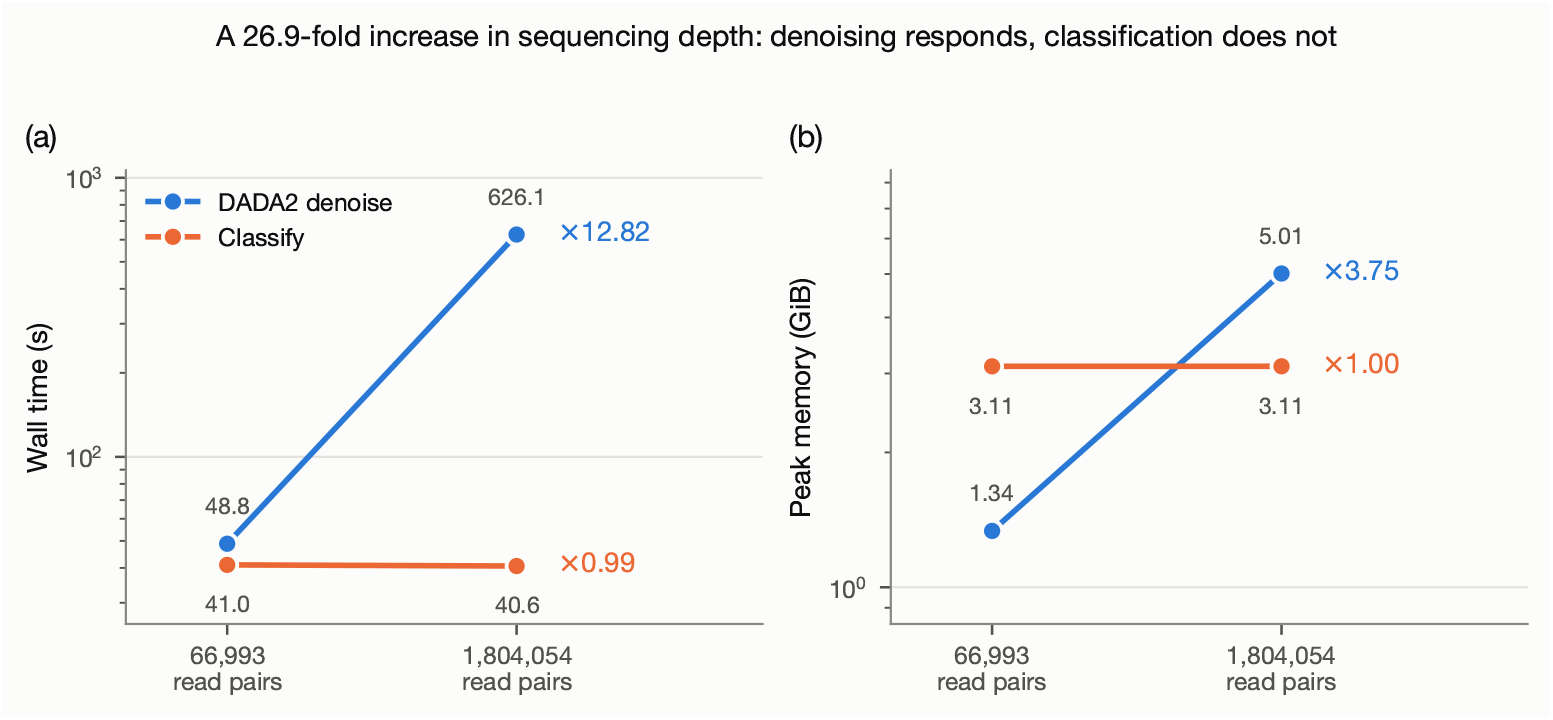
Sequencing depth drives denoising cost but not classification cost. Two nasopharyngeal samples from PRJEB33591, differing 26.9-fold in sequencing depth (66 993 versus 1 804 054 read pairs) but yielding only 54 versus 58 ASVs. Both stages run at four cores; points are means of three repetitions. **(a)** Wall time. **(b)** Peak resident set size. Note the logarithmic vertical axes. Ratios to the right of each series are deep/shallow.

Denoising responded strongly to depth, though sub-linearly: a 26.9-fold increase in reads produced a 12.8-fold increase in wall time, an empirical exponent of 0.78, and peak memory rose 3.8-fold. Classification did not respond appreciably. Wall time differed by 1% (41.02 s versus 40.64 s; Welch *t* = 1.18, *p* = 0.35) and peak memory was identical to three significant figures. With three repetitions this test has little power, so it bounds the effect without demonstrating invariance: the 0.383 s difference has a Welch 95% interval of [*−*0.87, +1.63] s (*ν* = 2.27), so the data exclude an effect larger than 4.0% of the stage and no more than that.

Framed as equivalence, two one-sided tests place the difference within a *±*4% margin (*p* = 0.025) but not within *±*3% (*p* = 0.055). That margin was not specified in advance and we do not present it as one: it is the bound this design supports, and it coincides with the resolution of Section 3.9 because both are governed by the same scatter. What makes 4% the right order of magnitude to care about is external to the statistics — classification takes 36 s to 47 s across every arm here, so 4% is under two seconds where scheduler requests are granted in whole minutes — and both statements would stop holding at larger query counts. The substantive comparison is the asymmetry: denoising changed by a factor of 12.8 where classification is bounded within a few per cent.

### 4.3 The same result within a single library

Because the two samples above are from different participants, the contrast was repeated within one library: ERR3444628 subsampled to six depths spanning 66 816 to 1 804 054 read pairs, holding composition exactly fixed (Table S6, Figure S2). Denoising rose 14.3-fold across the range, an empirical exponent of 0.817, closely matching the 0.78 of the between-participant contrast. Classification changed by 1%.

The ASV counts in Table S6 are a rarefaction curve for this library, rising with sampling effort without reaching an asymptote (Gotelli and Colwell, 2001; Willis, 2019), which shows the absence of a cost response is not simply richness saturating: ASV count rose from 21 to 58, a factor of 2.8, and classification time still did not move. Regressed on read count it gives *R*^2^ = 0.25, on ASV count *R*^2^ = 0.21, both describing the same 0.4 s of variation, of the order of the run-to-run standard deviation. Over the range these workloads occupy the per-ASV term is small enough relative to the fixed cost that a threefold change in richness is not resolvable.

### 4.4 Cost against richness across nine communities

Both stages were then measured on each of the nine samples individually (Figure S3). Denoising time follows read count with a power-law exponent of 0.75 (*R*^2^ = 0.98 on log axes); the three arms therefore place the exponent between 0.75 and 0.817 depending on which contrast is fitted, these being separate estimates of the same quantity rather than one value quoted inconsistently. The fit is carried substantially by the single deep sample: restricted to the eight between 44 604 and 82 257 reads it gives *R*^2^ = 0.67 and *r* = 0.82, which we report descriptively because those eight come from six participants, two contributing two sites each, so a nominal *p*-value would overstate the evidence.

Classification shows no detectable relationship with either predictor over this range: against ASV count *R*^2^ = 0.003, and total wall time varies by 0.46 s across communities spanning 5 to 70 ASVs. The extremes make the point more directly. The clonal effusion (ERR3444623; 5 ASVs) and the most diverse adenoid sample (ERR3444641; 70 ASVs, *H^′^* = 2.58) differ 14-fold in richness and are ecologically different kinds of community. They cost 40.81 s and 40.79 s to classify, a ratio of 1.000.

### 4.5 Classification is dominated by a fixed cost

The ASV sweep explains why: nearly all of the stage is paid before any sequence is classified. At one job, classifying a single sequence took 36.20 s and 218 sequences 37.27 s (Table S7, Figure 2); regressing wall time on ASV count gives an intercept of 36.25 s over this range and a slope of 5.21 ms per ASV, so at the largest set measured the fixed cost accounts for 97.0% of the stage.

**Figure 2.**
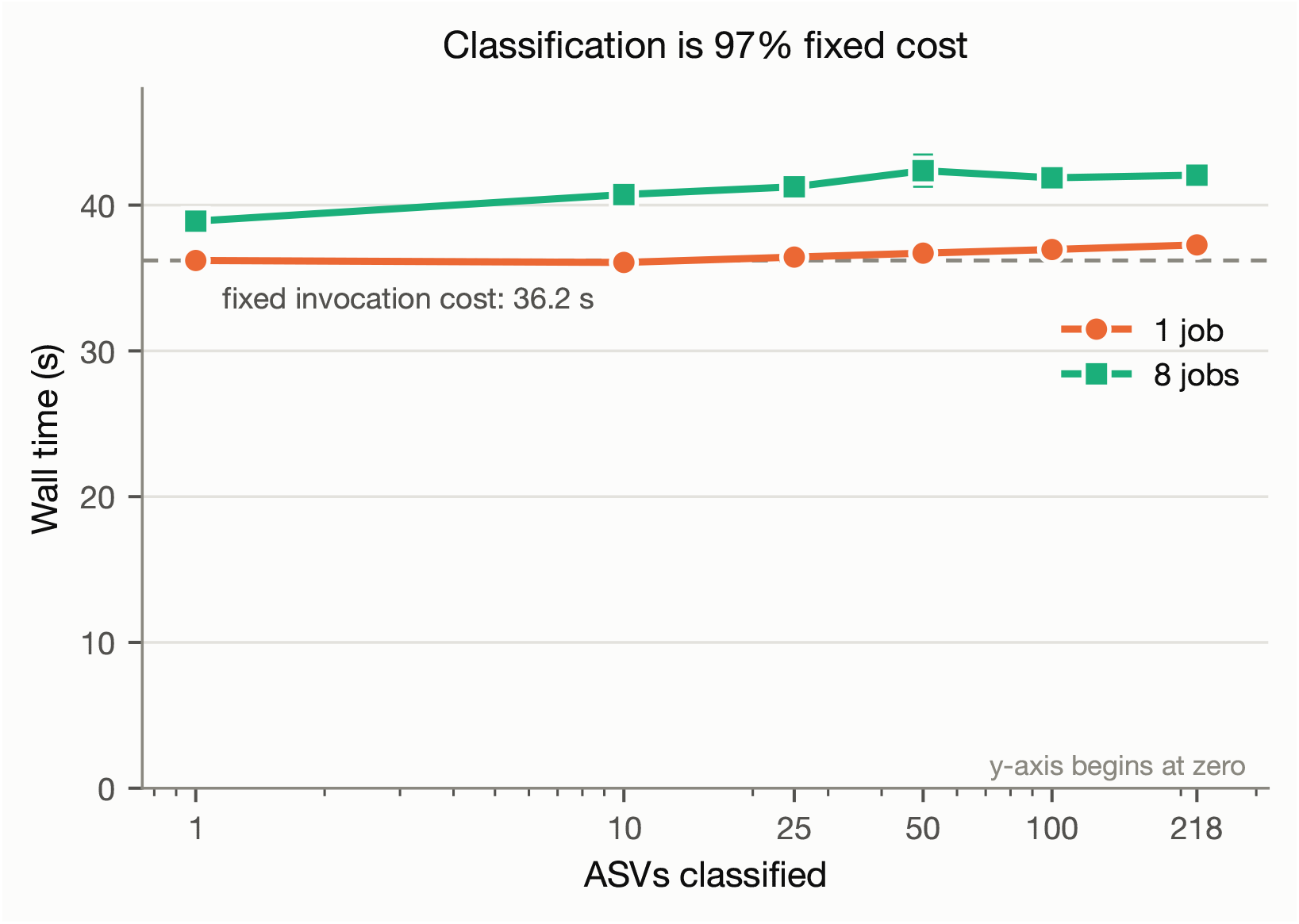
Classification cost is almost entirely fixed. Wall time for `classify-sklearn` against nested subsets of the 218 distinct representative sequences pooled across the nine samples, at one and eight jobs. Error bars are *±*1 SD over three repetitions; the dashed line marks the single-sequence time at one job, which is the fixed per-invocation cost of the stage. Eight jobs is slower than one at every set size measured.

We measured what that intercept consists of rather than inferring it, timing each candidate layer as a nested ladder, ten repetitions per layer in randomised order, so that consecutive rungs differ by one layer’s own cost (Table 4). Interpreter start-up and framework import account for 0.84 s, under 3%; one rung, `Artifact.load` of the reference, accounts for 19.20 s — and is not what its name implies. Since each rung is a longer prefix of the same command rather than a separable operation, we split it directly (Table 4, lower panel): importing the plugin namespace with no artifact touched costs 16.23 s, the `Artifact.load` of the 62 MiB reference a further 0.15 s, and unpickling it into a scikit-learn pipeline 2.04 s. Reading and deserialising the reference is therefore about 2.2 s, under 7% of the stage; it is not thereby absolved, since it reappears in the scoring call below and totals roughly 6.6 s, but it does not account for the rung it seemed to.

**Table 4.** What the fixed classification cost consists of. Each row adds one layer to the row above, so the right-hand column is that layer’s own cost. Ten repetitions per layer in randomised execution order; intervals are percentile bootstrap. Wall time is 100% CPU throughout. The `Artifact.load` row is not what its name suggests; the lower panel splits it directly, and reading the artifact is a small part of it. The 2.04 s pipeline view in the lower panel is the same quantity as the 2.11 s upper-panel row measured again, not an additional cost, and the lower panel’s absolute values come from a session that ran 7% faster (Section 3.9); the attribution is unaffected.

| Cumulative | Wall (s) | This layer | Share |
|---|---|---|---|
| Python interpreter start-up | 0.01 | 0.01 | 0.0% |
| + import qiime2 | 0.77 | 0.76 | 2.1% |
| + q2cli and the plugin manifest | 0.85 | 0.07 | 0.2% |
| + the feature-classifier plugin | 0.85 | 0.00 | 0.0% |
| + Artifact.load the reference | 20.04 | 19.20 | 53.0% |
| + materialise the sklearn pipeline | 22.15 | 2.11 | 5.8% |
| + the remainder of the invocation | 36.23 | 14.07 | 38.8% |
| *Splitting the Artifact.load rung directly (three repetitions, later session):* |  |  |  |
| import qiime2 | 0.78 | 0.78 | — |
| + instantiate the PluginManager | 17.01 | 16.23 | — |
| + Artifact.load the reference | 17.16 | 0.15 | — |
| + view it as a scikit-learn pipeline | 19.19 | 2.04 | — |

The dominant component is framework start-up of a particular kind. Resolving a semantic type makes QIIME 2 instantiate its `PluginManager`, which imports every installed plugin rather than the one being called: 25% of profile samples fall in `rpy2`, the R interface loaded on behalf of `q2-dada2`, and 6% in `llvmlite`, neither of which classification uses. The ladder’s own plugin rung missed this because `feature-classifier --help` reads a cached manifest where an invocation carrying artifact arguments does not.

For the work remaining after the pipeline exists we report the clock rather than the profile, which places 71% of in-action samples in `scipy.sparse`’s `matmul multivector` where direct timing puts the scoring call at 28% of the action. The action’s 16.1 s comprises a 1.95 s pipeline view, a 4.50 s scoring call and 9.65 s of other work. A normal run pays that view once, inside the action: the phase-timed invocation enters the action having only opened the artifact handle (0.008 s), and the explicit view rung of Table 4 prices the same operation rather than an additional one. The scoring call costs the same every time (4.503 s, 4.498 s, 4.502 s and 4.501 s over four consecutive calls on one query), so it is per-call work and not first-call setup, and barely moves with query count: all 218 sequences cost 4.92 s against 4.50 s for one, 1.9 ms per additional sequence, agreeing with the 2.497 ms slope fitted independently in Section 4.8.

A per-call cost that does not scale with queries is what a fixed cost is, and the reference is what sets it: the fitted likelihood matrix is 10 785 classes by 8 192 features in `float64`, 674 MiB resident against a 62 MiB artifact on disk, traversed once per call. We do not claim the 4.50 s is the arithmetic itself, which for a sparse single-query vector would be far cheaper; the shape of the cost says the array is traversed, not how efficiently. The remaining 9.65 s is QIIME 2’s own machinery, including provenance capture and the 2.65 GiB of scratch each invocation writes, which we have not decomposed further. The command-line interface is not implicated: 33.60 s through the Python API against 33.48 s through `q2cli`.

The attribution is worth stating precisely, because the scratch volume invites a different one. Each classification writes 2.65 GiB to the filesystem, essentially independent of ASV count and some forty times the size of the reference artifact on disk (denoising, by comparison, writes 33 MiB), which suggested the artifact was being unpacked to scratch on every invocation and that the cost was therefore storage-dependent and perhaps avoidable through caching. It is not. Placing the classifier on `tmpfs` rather than NVMe changed the full classification by 0.00 s (95% CI *−*0.07 s to 0.07 s) and the artifact load by 0.05 s (*−*0.05 s to 0.15 s), and 99.9% of the 36.23 s total is CPU time. The stage is compute-bound, and mostly in module import, not waiting on storage. The scratch write is real but is not the bottleneck. Because both `tmpfs` arms ran with a warm page cache, we also measured the cold case directly, dropping the page cache immediately before each of five classifications and pairing each with a warm run that followed it. Cold runs cost 37.99 s against 36.08 s warm, a paired difference of 1.90 s (95% CI 1.83 s to 1.97 s), or 5.3% of the stage; the cold runs read 501 MiB from storage and were still 96% CPU.

A second cold-cache series, run later to locate that penalty, attributes most of it to the import path rather than to the reference. Dropping the cache before an import-only run, which touches no artifact, raised it from 17.02 s to 19.10 s; dropping it before a full classification raised that from 33.30 s to 35.83 s. Of the 2.53 s cold penalty, 2.08 s, or 82%, is paid importing modules, and recovery is immediate, the first classification after a drop costing 35.56 s and the next five 33.66 s on average. Storage therefore contributes at most about two seconds of the 36 s fixed cost even when nothing is cached, so this cost scales with processor speed rather than storage and a faster disk will not reduce it. A smaller reference would help, though less than the plugin set: the reference accounts for about 6.6 s of the stage, and we measured one reference, so we cannot say how either of its terms scales with matrix size. The larger lever is the installed plugin set, which is both bigger and, as Section 4.6 shows, free to pull.

Peak memory was likewise almost unaffected, rising 18 MiB across the whole range, from 3.109 GiB at one sequence to 3.127 GiB at 218.

The per-ASV term sits close to the measurement noise: total wall time varies by 1.2 s across a 218-fold change in input size against a run-to-run standard deviation of roughly 0.1 s, and the linear fit is correspondingly modest (*R*^2^ = 0.86), carried largely by the largest point (at *k* = 218 the leverage is *h* = 0.84 and Cook’s distance (Cook, 1977) is 6.8, where no other point exceeds 0.4; removing it moves the slope from 5.21 to 8.65 ms per ASV). We regard the flatness as the substantive finding rather than a limitation of the fit: over the range of ASV counts these workloads actually occupy, classification time is effectively constant. Propagating the slope’s 95% interval (2.33 ms to 8.10 ms per ASV), the point at which per-ASV work would equal the fixed cost lies between roughly 4 500 and 15 600 — a three-fold uncertainty, 32 times beyond the largest set measured, which indicates an order of magnitude and no more. That is what motivated the extended query sweep of Section 4.8, which measures the range directly instead of inferring it.

### 4.6 Nearly half the fixed cost is removable

If the dominant component is the framework importing plugins the job does not use, removing them should remove the cost. We built a second environment from the same channels pinned to the same versions, requesting the framework, `q2cli`, `q2-types` and `q2-feature-classifier` and nothing else. Dependency resolution adds `q2-feature-table`, `q2-quality-control` and `q2-taxa`, giving 5 plugins against 27, 377 packages against 683, and no `rpy2`, since `q2-dada2` is absent. Ten single-sequence classifications were run in each environment, interleaved in randomised order within one session so that the session term of Section 3.9 and any NUMA placement effect fall on both arms equally, with five import-only runs each.

The saving is large and it is free (Table 5). Classification falls from 33.24 s to 18.75 s, a reduction of 43.6% (95% CI *−*14.61 s to *−*14.38 s), and the import phase alone from 17.08 s to 2.69 s, a reduction of 14.38 s. That the two agree to 0.11 s is the point: the entire saving is import, and nothing else about the invocation changed. Every run in both environments produced a byte-identical taxonomy table (SHA-256 `95875c93ee05a864…`), so the 43.6% is not bought with any change in the result. Peak memory falls from 3.10 GiB to 2.22 GiB, 28.3%, which matters more for scheduling than the wall time does because Section 5 sizes the memory request by the classification stage: the request falls from 7 GiB to 5 GiB.

**Table 5.**
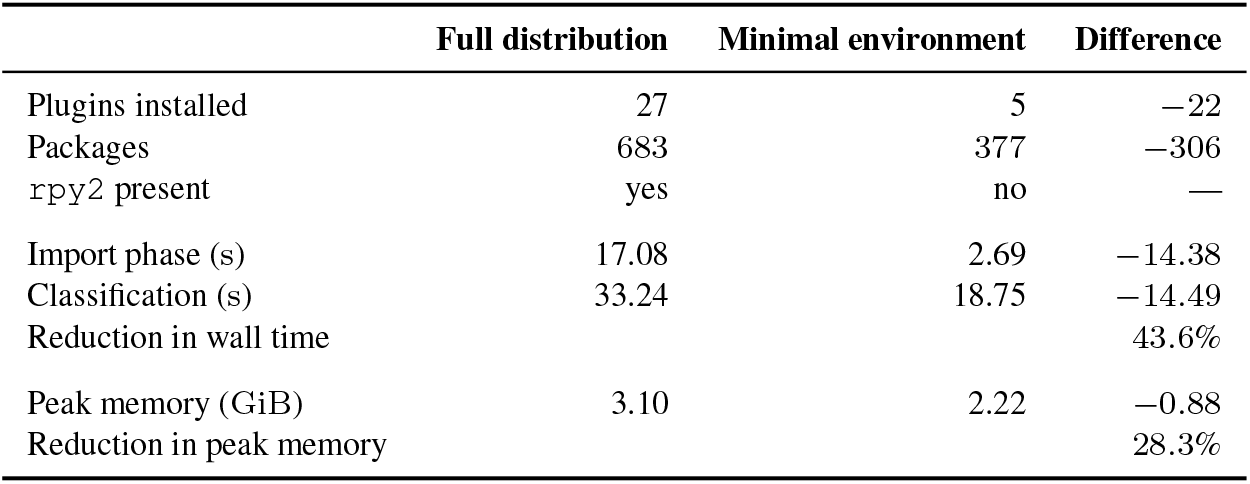
The same classification in the full distribution and in a minimal environment, interleaved in randomised order within one session. Intervals are percentile bootstrap on the difference of means. The taxonomy table was byte-identical in all 20 runs.

The environment is reconstructible from the recipe and not only from the export. Building it afresh from `environments/qiime2-lean.yml` on a machine where it did not exist, eighteen days after the measurements above, returned the same 5 plugins, the same byte-identical taxonomy table and the same 2.22 GiB peak, with every pinned version held. In the interval 51 transitive dependencies had moved to newer patch releases and two that the stage does not use, `pylatexenc` and `six`, had dropped out of the solve. The container image built from the same recipe returns the same digest again on a different machine and processor architecture. The result is reproducible; the exact package set is not, which is why the provenance export is kept alongside the recipe rather than replaced by it.

Two caveats belong with this. The minimal environment cannot run the pipeline: denoising needs `q2-dada2`, which is what brings in the R runtime, so the saving is available to classification run as a separate step rather than to a single environment doing everything. That is not a serious obstacle on a cluster, where stages are already separate jobs, but it does mean the recommendation is about job design and not merely about installation. And the figure is specific to this distribution and this plugin set; what transfers is the method of measuring it, not the 14.49 s.

### 4.7 Thread-level parallelism is ineffective, and harmful for classification

Neither stage converts additional cores into wall-time savings, and for one the effect is negative (Table S8, Figure 3). Denoising speedup peaked at 1.184*×* near eight threads and declined to 1.167*×* at 40: going from 1 to 40 threads bought 14% of wall time at 2.3 times the CPU, with voluntary context switches rising from 836 to 26 751. Classification was slower than serial at every setting above one job, taking 1.28 times as long at 40 jobs while consuming 10.5 times the CPU; the stage uses `joblib` (Pedregosa et al., 2011) with its process-based `loky` backend, and with only tens of sequences to distribute, worker startup and result collection exceed the work distributed.

**Figure 3.**
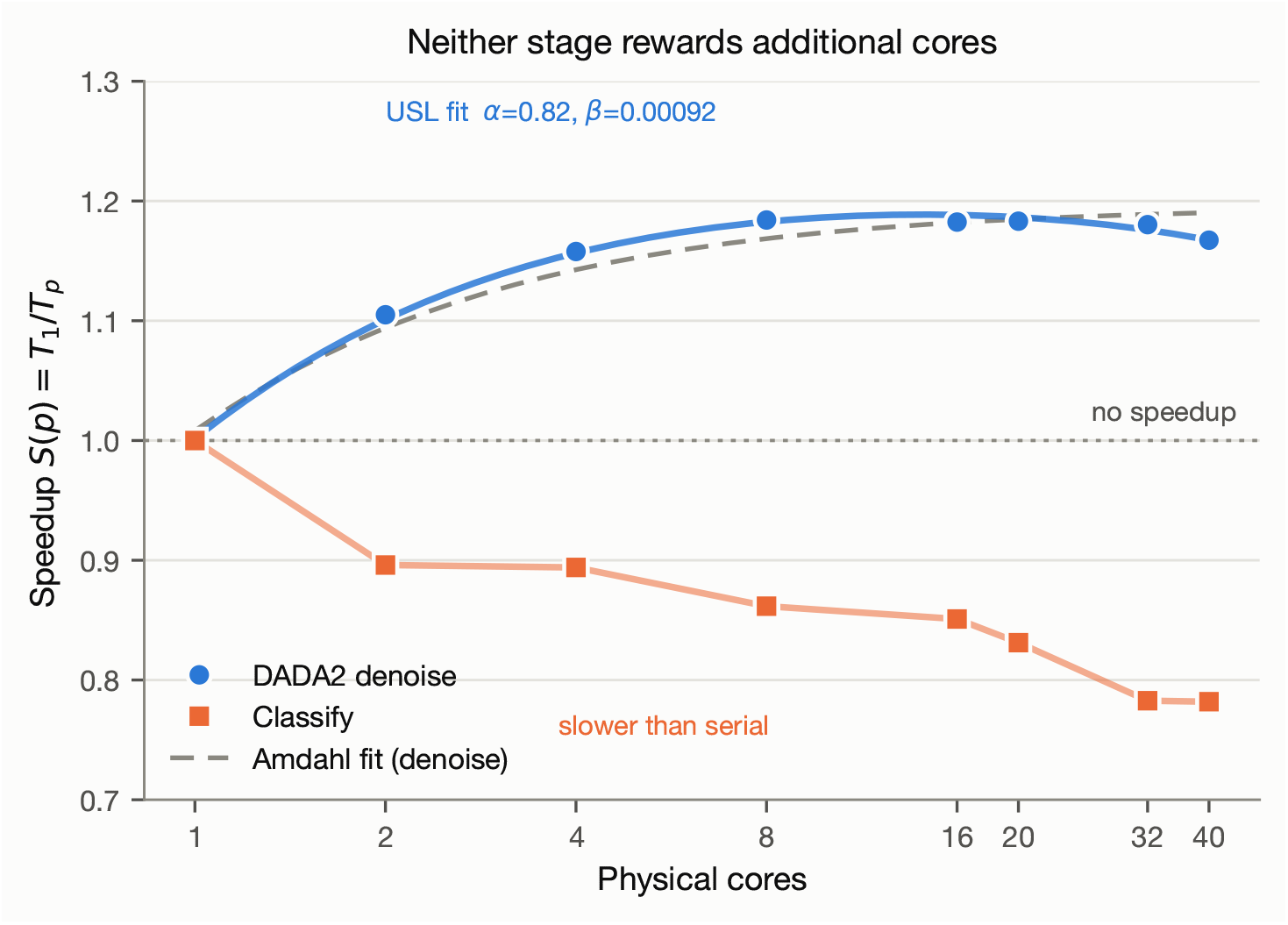
Neither stage rewards additional cores. Strong-scaling speedup *S*(*p*) = *T*_1_*/T_p_* against physical core count, means of three repetitions. Denoising (ERR3444605, 66 993 read pairs) peaks at 1.184*×* near eight threads and declines; the solid curve is the fitted Universal Scalability Law and the dashed grey curve the fitted Amdahl’s Law, which is monotonic and so cannot reproduce the decline. Classification (54-sequence ASV set) is below serial performance at every setting above one job. The classify series is drawn with a connecting line only; no scaling model is fitted to it.

Those two ratios also bound the frequency confound of Section 3.3: a clock change scales wall and CPU time by the same factor and cannot create CPU seconds that were not there, so a 1.28-fold rise in wall time against a 10.5-fold rise in CPU means the great majority of the additional CPU time is work introduced by parallelism. For denoising the confound runs the other way, wall time *falling* to 0.86 of serial at 40 threads, so the matched-frequency ceiling would be somewhat higher rather than lower. The ASV sweep shows the same pattern independently: at every set size eight jobs were slower than one, by 1.13*×* at 218 sequences.

Fitting the denoising curve, Amdahl’s Law gave *f* = 0.843 with *R*^2^ = 0.961, while the Universal Scalability Law gave *α* = 0.818, *β* = 0.000 916 with *R*^2^ = 0.994 and an optimum of *p^∗^*= 14 cores.

Resampling the replicates (5 000 percentile bootstrap draws) gives *f* a 95% interval of 0.839 to 0.847, *α* 0.812 to 0.824, *β* 0.000 712 to 0.001 099 and *p^∗^* 13 to 16 cores. Those intervals describe repeatability under the fitted model, not its adequacy: the *measured* peak lies at eight threads and speedup between eight and 32 varies by less than 0.4%, so *p^∗^* locates a broad flat region rather than a sharp maximum. Penalising the Universal Scalability Law’s extra parameter it remains favoured (adjusted *R*^2^ 0.996 versus 0.951; ΔAIC*_c_* = *−*12.1, computed on the 8 per-core means). The difference matters for prediction: Amdahl’s Law is monotonic and continues to predict improvement out to 40 cores, where the stage is measurably slower, so a predictor built on Equation 1 would under-predict wall time on the declining branch — the situation a user creates by over-requesting cores.

### 4.8 Where the fixed cost stops dominating

The results above are confined to the range a single sample occupies, at most 218 sequences, and per-query work must eventually dominate any fixed cost. We built a larger query set from ERR3444628 by taking R1 reads before denoising, discarding those with ambiguous bases, trimming to 250 bp and dereplicating, yielding 60 000 distinct sequences; fixing length leaves query count as the only variable. Subsets from 1 to 60 000 were run at one job and at eight, three repetitions each, the ladder extended past 10 000 so that the crossover is bracketed by measurement rather than reached by extrapolation. Where the two query sets overlap (1, 10, 50, 100 and 218) they agree to 0.65% on average and at most 1.8%, in both directions rather than as a systematic offset (Table S9), so the ladder above 218 continues the same curve.

At one job the line *T* (*k*) = *a* + *bk* describes the data closely over the full ladder (*R*^2^ = 0.999 5), with *a* = 36.36 s and *b* = 2.497 ms per query. That intercept is within 0.3% of the 36.25 s fitted independently over 1 to 218 ASVs, from a different query set spanning a range 275 times narrower, and within 0.4% of the 36.23 s measured directly by the decomposition. Fixed and accumulated per-query costs are equal at *k^∗^* = *a/b* = 14 560 queries (95% CI 14 437 to 14 714, Figure 4a) — the fixed-to-variable dominance crossover, a different quantity from the parallel-efficiency crossover below. It does not depend on the fitted line at all: a single query costs 36.49 s, so the fixed cost is matched at 72.97 s, and the measured points at 15 000 and 20 000 queries (71.86 s and 83.91 s) place the crossing near 15 500 by interpolation, 6% from the fitted estimate. Two earlier extrapolations can be checked against this: from 1 to 10 000 queries, 15 248; and from 1 to 218 ASVs, an interval of 4 500 to 15 600, which contains it. For scale, the pooled union across all nine samples is 218 distinct sequences, roughly 70-fold below *k^∗^*.

**Figure 4.**
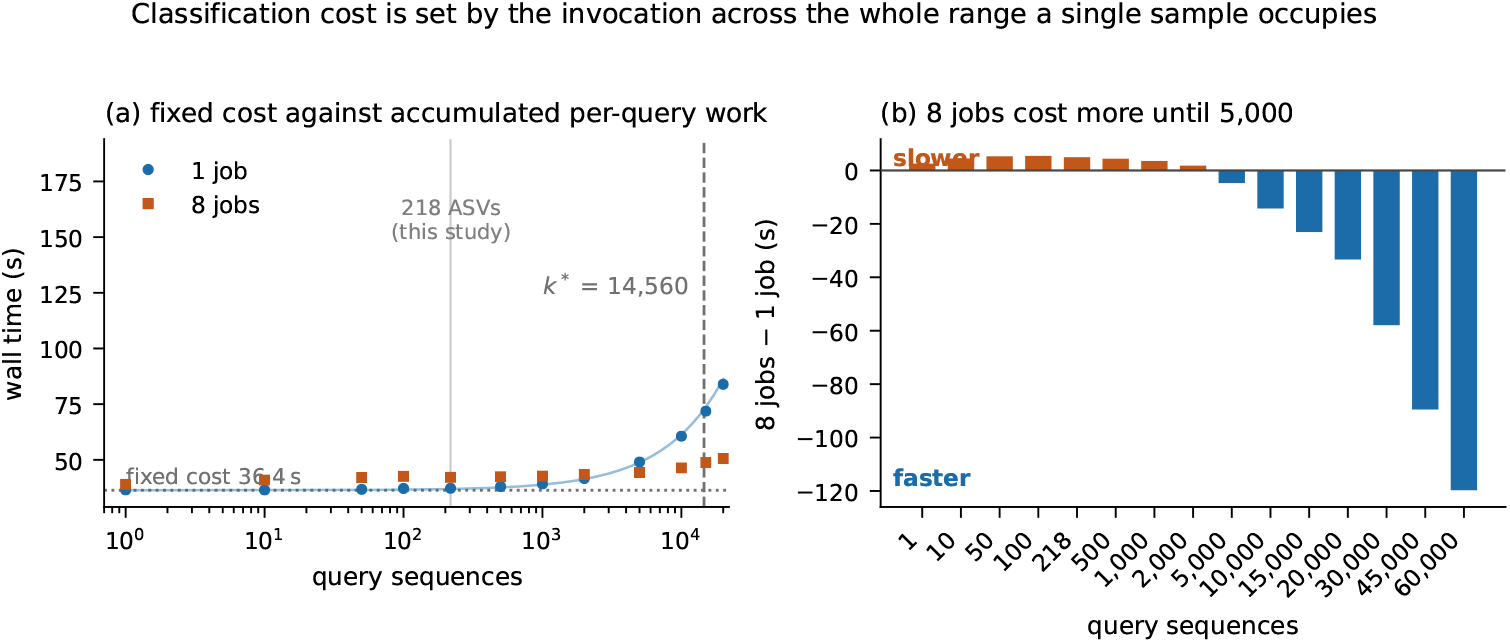
Classification cost is set by the invocation across the whole range a single sample occupies. **(a)** Wall time against query count at one job and eight, means of three repetitions, error bars one standard error, logarithmic abscissa. The dotted line is the fitted fixed cost *a* = 36.36 s and the dashed line the crossover *k^∗^* = *a/b* = 14 560 at which accumulated per-query work equals it. The 218 sequences this cohort produces are marked. **(b)** The difference between the two settings. Positive values are query counts at which eight jobs take longer than one; the penalty peaks at 5.46 s at 100 queries and persists to 2 000. Queries are 250 bp dereplicated reads from ERR3444628; at the five query counts where this sweep overlaps the ASV sweep, the two agree to within 1.8%.

At eight jobs a straight line does not describe the data (*R*^2^ = 0.710): cost climbs 3.60 s over the first 99 queries, is flat to approximately 2 000, and only then becomes linear, so we report no crossover from it. Eight jobs are slower than one at every count up to 2 000, by 1.80 s to 5.46 s (Figure 4b), the penalty largest at 100 queries where 5.46 s is 14.7% added to a 37.17 s job; they first win beyond replicate scatter at 5 000 and reach 1.31*×* at 10 000. That threshold is the parallel-efficiency crossover and should not be conflated with *k^∗^*: one is where a second configuration overtakes the first, the other where two terms of a single model become equal, and they differ threefold here. The instruction is not that additional jobs stop helping below either threshold, but that requesting them makes the job measurably worse.

### 4.9 Output is unchanged by thread count

Changing the number of threads requested is only safe if the biological result does not change with it, and DADA2 estimates its error model from a subsample of reads, so this has to be shown rather than assumed. Denoising ERR3444605 at 1, 4 and 40 threads produced 54 ASVs in every case with identical sorted representative-sequence sets (SHA-256 `c759115c59be27c5…`), and classifying those sequences at the same three settings produced 54 assignments identical in both feature and taxon (SHA-256 `11b2812c…`). The second check is the one that matters: identical sequences would not by themselves guarantee identical taxonomy, and it is the taxonomy a study reports. Core count is therefore a pure performance parameter here. The scope of the check is given in Section 6.1.

### 4.10 Memory

Denoising memory was insensitive to thread count (1.31 GiB at one thread to 1.36 GiB at 40) but responded to input size. Regressing peak RSS on read count across the within-sample depth series, in which thread count is held fixed at four (Figure S4a), gave

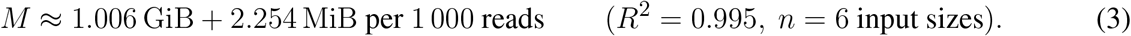

The intercept has a 95% interval of 0.824 GiB to 1.187 GiB and the slope 2.037 MiB to 2.472 MiB per 1 000 reads, the intercept being the quantity to treat with most caution when transferring the model since it extrapolates to zero reads. The weak-scaling series is the more obvious source for this fit and is the wrong one: it scales read count *in proportion to* thread count by construction, so the two predictors are perfectly correlated (*r* = 1.000) and up to 40% of the apparent read effect is attributable to thread count. Nor can a memory model be fitted from a strong-scaling series alone, since read count does not vary within it. Classification memory (Figure S4b) was constant at 3.11 GiB across every condition measured, so it is a property of the invocation and not of the data.

### 4.11 Predictor accuracy on within-cohort held-out samples

The models above are only useful if they predict inputs they were not fitted to. Time and memory were calibrated on the within-sample depth series alone, giving *T* (*n*) = 37.51 s *×* (*n/*66 816)^0.817^, *M* (*n*) = 1.006 GiB + 2.254 MiB per 1 000 reads, and a per-sample classification time of 40.93 s at four jobs — the mean of that series’ own classification runs, not the fixed per-invocation cost, which is 36.2 s at one job, the difference being the penalty for requesting four. They were then used to predict the per-sample measurements of eight other libraries from different participants and body sites, collected in a separate arm (Table S10; ERR3444628 is excluded because the depth series was built from it). The validation is within-cohort: all eight libraries come from the same study, the same sequencing platform and the same preparation protocol as the calibration library, so what is tested here is transfer across communities and body sites, not across platforms, studies or biomes. Transfer across those remains untested and is treated as a limitation in Section 6.1.

The three models do not transfer equally well. The fixed classification cost predicts the within-cohort held-out samples to within 0.7%, mean 0.34% — the strongest evidence in this work that the quantity is a property of the invocation rather than of any input, eight communities spanning 5 to 70 ASVs all costing what one subsampled library cost. Denoising peak memory predicts to within 13.7%, mean 7.9%.

Denoising wall time is the exception and fails in the dangerous direction, under-predicting every within-cohort held-out sample by 29% on average and 40.9% at worst. Two causes contribute. The power law does not fit its own calibration series at the shallow end — at 66 816 reads it returns 37.51 s against a measured 43.67 s, *−*14.1% — and its residuals across the series (*−*14.1, +7.4, +9.2, +9.7, +1.9, *−*11.2%) show the data are not a pure power law and the fit is worst exactly where the held-out samples lie, so roughly half the 29% mean error is misfit rather than anything to do with the libraries being different. The rest is genuine between-library variation, which a calibration built by subsampling one library cannot see because DADA2’s work per read depends on the error profile of the run. An under-predicted `--time` is what kills a job, which is why the worked example of Section 5.1 doubles the estimate, and why a reader calibrating for production should fit the exponent across the libraries they intend to run rather than by subsampling one of them.

## 5 RESOURCE RECOMMENDATIONS

The measurements support five recommendations. All are drawn from warm-cache measurements on a single node against one reference artifact, and the constants should be remeasured before being relied on elsewhere.

1. **Size denoising by read count; calibrate classification against the installation rather than against the data.** Denoising follows reads^0.817^ in time and Equation 3 in memory, while classification is set by a fixed per-invocation cost that read count does not enter. That constant belongs to the installation and the reference together: about 16 s is the framework importing its plugins and about 6.6 s the reference. Measure it once for the environment and reference in use.
2. **Request one job for classification below roughly** 5 000 **query sequences.** Every setting above one job was slower across 1–218 sequences, and this is not a small-input artefact: eight jobs remain slower out to 2 000 queries, by as much as 14.7% at 100, first win at 5 000 and reach 1.31*×* at 10 000.
3. **Request few cores for denoising.** Two cores reach 1.105*×*, 93% of the best speedup observed at any core count, at 55% efficiency; four reach 98% of it. Beyond eight, additional cores cost CPU and return nothing.
4. **Obtain throughput from sample-level parallelism.** Since per-sample threading saturates below 1.2*×*, concurrent low-core jobs dominate, and we measured this rather than projecting it. Running *N* identical single-core classifications simultaneously on the 40-core node, throughput rose monotonically to 21.1*×* the single-job rate at *N* = 40 with no failures (Table S11), per-job wall time rising from 37.59 s alone to 68.31 s under full load. That baseline is one single-core job on an idle node, not the threading one; against four jobs per sample in four waves (4 *×* 41.03 s = 164 s), 40 concurrent single-core jobs complete the same work in 71.40 s, an advantage of 2.30*×*, both sides being classification at the same query count. Memory scales linearly with headroom to spare: 115.5 GiB aggregate at forty jobs, 46% of the 251 GiB installed.
5. **Run classification in a minimal environment.** Nearly half the fixed cost is the framework importing plugins the job never calls. An environment holding only the framework, `q2cli`, `q2-types` and `q2-feature-classifier` classifies the same sequence in 18.75 s against 33.24 s, a reduction of 43.6% with a byte-identical taxonomy table (Section 4.6), and peak memory falls from 3.10 GiB to 2.22 GiB. Denoising still needs the full distribution, so this applies to classification run as its own job, which on a cluster it usually is. It is the largest single saving available here, and unlike the others it costs nothing in allocation or in scheduling.

### 5.1 Worked example: from measurements to a scheduler request

Take a sample of 500 000 read pairs, denoised at four cores then classified. Anchoring on the measured 43.67 s at 66 816 reads, *T*_denoise_ *≈* 43.67 s *×* (500 000*/*66 816)^0.817^ *≈* 223 s. Classification contributes its fixed cost: about 37 s at one job, which is what Recommendation 2 calls for at this query count, rather than the 41 s four jobs would cost, for a total near 260 s. Memory is the larger of the two stages rather than their sum, since they run in sequence: Equation 3 gives denoising 2.11 GiB against classification’s fixed 3.11 GiB, so the request is sized by classification even though denoising takes five times as long. Doubling both for margin:

~~~
#SBATCH --time=00:10:00
#SBATCH --mem=7G
#SBATCH --cpus-per-task=4
~~~

Splitting classification into its own job in a minimal environment (Recommendation 5) would ask `--mem=5G` and about 19 s rather than 37 s for that step. The figures above are for a single job running both stages in the full distribution.

Under a workflow manager the split costs nothing in pipeline structure, because both Nextflow and Snakemake attach an environment or a container to the individual process rather than to the run. In Nextflow it is a `container` or `conda` directive on the classification process alone, selected per process through `withName`; in Snakemake it is a `container:` or `conda:` key on that rule alone. Every other step continues to use the full distribution, which it needs. The denoising step cannot use the lean image: `q2-dada2` is absent from it by construction, which is where most of the 43.6% comes from.

Equation 3 is an interpolation guide and should be read as one. It was fitted to six input sizes from a single library spanning 66 816 to 1 804 054 read pairs, and it is reliable only within that range: the worked example above sits inside it, but a library an order of magnitude deeper or shallower lies outside the calibration and the intercept in particular extrapolates badly, as Section 4.10 notes. A reader provisioning for production should refit the two coefficients on the libraries they intend to run rather than adopt the values printed here. The same applies with more force to the denoising time exponent, which Section 4.11 shows under-predicting by 29% on average even within this cohort.

The margin is asymmetric: an underestimate kills the job, an overestimate wastes memory no other job can use. The heuristic this replaces, sizing classification in proportion to read count as our own earlier SLURM templates did, would scale the request by 7.5 and ask for some 23 GiB where the stage uses 3.11 GiB regardless of depth.

### 5.2 Which unit to size against

The recommendations are stated per invocation, and the quantity that matters is the number of *distinct* sequences presented in one call — neither the number of samples nor the number of reads. Classifying one sample at a time presents that sample’s ASVs alone, so the fixed cost is paid once per sample and a 96-sample study pays it 96 times. Classifying a batch presents the union, which grows sub-linearly with sample count because samples share taxa: our nine carry 374 representative sequences but only 218 distinct ones. That the fixed cost is then paid once is a stronger argument for batching than any saving in per-query work. Union size is community-dependent — hundreds to thousands of distinct sequences are ordinary for soil or sediment (Thompson et al., 2017) — and above roughly 5 000 queries the parallel recommendation inverts while above roughly 15 000 the per-query term overtakes the fixed cost, both thresholds being properties of this node and reference. Table S12 maps the three cases onto the allocation each implies.

## 6 DISCUSSION

A 16S pipeline does not have a single input size. Denoising is a function of read count; classification is a function of the installation it runs in and of the reference it loads, with a weak dependence on query count, and over the range these workloads occupy it is dominated by a cost that varies with neither. A middle-ear effusion carrying a single organism and an adenoid community of seventy variants are about as different as two samples from one study can be, and they cost the same to classify to three significant figures. A 27-fold increase in depth changed classification cost by 1%, whether from comparing two participants or from subsampling one library, so a heuristic that scales the classification request with read count over-provisions by more than an order of magnitude on the deeper sample here. The within-sample series also separates two explanations the between-sample contrast leaves entangled: richness did not saturate, yet cost still did not move, so the mechanism is not that deeper sequencing stops yielding new ASVs but that the per-ASV term is negligible against the fixed cost — a stronger and less conditional statement than saturation would support.

Two further results are more surprising. Denoising returns almost nothing for additional threads, with a measured ceiling of 1.18*×* on a single sample, and classification is actively degraded by parallelism, running 28% slower at 40 jobs than at one; because representative sequences were identical at every thread count tested, reducing the request costs nothing. Neither result extrapolates to multi-sample artifacts, and Thompson et al. (2022) provide the comparison we cannot: over 96 samples, denoising fell 3.3-fold from one core to eight against our single-sample 1.18*×*, since DADA2 distributes across samples.

For this classifier the relevant comparison is Bokulich et al. (2018), who fitted runtime for the same implementation over query sets up to 10 000 sequences, reported 23 ms per query and set the intercept aside as negligible. Our measurements are of that intercept, in the regime where it has not yet become negligible; Section 4.8 measures the transition at 14 560 queries, and that it sits so high is the substantive point. Our slope, 2.497 ms per query, is an order of magnitude below the 2018 figure, but the two come from different machines eight years apart and nothing here isolates a cause — itself an argument for calibrating locally rather than reading a constant off a published table.

That the Universal Scalability Law fitted where Amdahl’s Law did not is worth noting: Amdahl’s Law is monotonic by construction and cannot express a workload that degrades, nor can Gustafson’s reformulation (Gustafson, 1988), and the limits of the two-parameter form under multicore synchronisation have been noted before (Hill and Marty, 2008). At 28% degradation a predictor built on it would under-predict wall time in exactly the configurations users request.

Independent support comes from Guéguen et al. (2026), who, benchmarking 38 combinations of algorithms, classifiers and databases for accuracy, report in passing that sequence resolution rather than taxonomic assignment was their CPU bottleneck. They attribute this to small databases and few variants; our measurements indicate a different mechanism, since classification shows no detectable response to variant count across the range a single sample occupies. That the same asymmetry appears on different hardware, tools and cohort suggests it is a property of the workflow rather than of our installation.

Resource prediction is studied, though almost entirely outside bioinformatics (Rodrigues et al., 2016; Tanash et al., 2019; Gaussier et al., 2015). That work treats the job as opaque and learns from scheduler logs; the present work opens one workload and measures which input dimension each stage responds to. A log-trained predictor would learn the same fixed cost eventually, as a constant it could not account for; measuring it directly says what the constant is made of and when it stops holding. These results likewise complement workflow managers (Di Tommaso et al., 2017; Kö ster and Rahmann, 2012), which decide what runs and in what order where the measurements here inform how much to request.

### 6.1 Limitations

#### One study, one platform

All samples come from PRJEB33591 and span 5 to 70 ASVs at comparable depth, plus one library subsampled across a 27-fold depth range — deliberate, since it removes batch and protocol confounders from a study whose dependent variable is ASV count, at the cost of a narrow richness range drawn from upper-respiratory communities. We attempted to include rhizosphere samples but excluded them on finding their reads retained PCR primers. The sweep of Section 4.8 extends the measured *cost* range to 60 000 queries and so does not depend on that richness, but its queries are dereplicated reads rather than ASVs from a genuinely rich community.

#### Single node, single hardware configuration

Alternative core counts and distributed filesystems are not examined, and NUMA effects only to the extent of showing that a cross-node placement costs 14.4%; we did not bind, and the measurements reported here were taken unbound. We were unable to obtain access to a materially different platform, so the transfer of these relationships across hardware remains untested, and the dimensionless relationships of Section 3.5 are stated as expectations the released protocol is designed to test locally. The one piece of indirect evidence is internal and speaks to stability on this node rather than transfer off it: the fixed classification cost was determined six independent ways, including a randomised replication eight weeks later, and agrees to 1.2% (Table 3).

#### Between-session variation in classification is not attributed

The same single-sequence classification measured 36.05 s in one session and 32.98 s to 33.47 s in another on the same idle machine, while denoising reproduced to 0.55%. Of six candidate causes five are excluded; the survivor, NUMA placement, we cannot establish retrospectively, so the effect is reported as real and unattributed and binding is now recommended as a control. Absolute classification constants here carry a session term of this order in addition to the quoted run-to-run scatter.

#### Replication covers the two scaling arms, not every arm

The remaining arms retain three sequential repetitions and so resolve the large effects reported and not small ones.

#### Warm page cache, single-sample denoising, one determinism dataset

Timings are warm-cache; the cold penalty was measured for classification (1.90 s) but not for denoising. DADA2 parallelises across samples as well as within them, so the scaling here, for single-sample artifacts, should not be extrapolated to multi-sample denoising: Thompson et al. (2022) measured a 3.3-fold speedup from one to eight cores on a 96-sample artifact against our 1.18-fold ceiling. Determinism across thread counts held, as expected, but was verified for one sample on one QIIME 2 release.

#### Concurrent throughput was measured on one node only

under genuine concurrent load rather than projected, but with one local filesystem; a cluster placing many such jobs against shared or networked storage may contend where this node did not.

#### Classifier-specific constants

The 36.25 s fixed cost and 3.11 GiB footprint belong to the 62 MiB artifact of Section 2.4, and *k^∗^* = 14 560 is a ratio of two such constants. We recommend calibrating once against the environment in use, but exercised that recommendation at one point rather than showing how the constants move when the installation changes. The decomposition reorders which second experiment matters most: a second reference artifact would probe the 6.6 s the reference accounts for, where varying the installed plugin set probes the 16 s one, which is why we tested that instead. How either reference term scales — the 2.2 s to read it or the 4.50 s to traverse its matrix — remains unmeasured. The qualitative result should transfer; the constants should be recalibrated locally, for the installation as well as for the reference and the hardware.

### 6.2 Conclusion

Denoising and taxonomic classification in a QIIME 2 amplicon pipeline consume different inputs and should be resourced differently. Everything below concerns `classify-sklearn` with one locally trained Naïve Bayes reference; we did not benchmark a second reference or an alignment-based classifier. Across a 26.9-fold range of sequencing depth, denoising wall time rose 12.8-fold while classification changed by 1% and its peak memory by 0.02%, and at 218 ASVs 97.0% of classification time is the fixed per-invocation component.

Decomposing that component locates it in an unexpected place: 16.23 s is QIIME 2 instantiating its plugin manager, which imports every installed plugin including an R runtime classification never uses, while the reference accounts for about 6.6 s — 2.2 s to read and deserialise it and 4.50 s in a scoring call set by its 674 MiB likelihood matrix rather than by the queries — and emptying the page cache adds only 1.90 s. It is removable: classifying in an environment holding only the plugins the step requires takes 18.75 s against 33.24 s and returns a byte-identical taxonomy, a 43.6% saving for a change of environment alone. Extending the query ladder to 60 000 sequences places the point at which per-query work overtakes the fixed cost at 14 560 queries, two orders of magnitude above a single sample.

Neither stage benefits appreciably from thread-level parallelism and classification is degraded by it, though running samples concurrently raises throughput 21.1-fold at 40 jobs against a single job alone, and 2.30-fold against the same work done four cores at a time. Across a cohort whose richness varies 14-fold for biological reasons, classification cost varied by 0.46 s between the cheapest and dearest of the nine samples and by 0.05% between the two extremes of richness, and neither the representative sequences nor their taxonomic assignments changed with thread count.

For practitioners: size denoising by depth; calibrate classification once against the software environment and reference in use rather than scaling it with sequencing depth; run it in an environment holding only the plugins it needs; request one job for classification at these input sizes and few cores for denoising; and take throughput from running samples concurrently instead of threading any single sample. The biological output is unchanged across the thread counts tested, so these reductions cost nothing in what the analysis reports. The instrumentation, data and analysis code are released so that these constants can be recalibrated for other installations, references and hardware.

## Supporting information

Data Sheet 1

Data Sheet 2

Data Sheet 3

Data Sheet 4

Data Sheet 5

Data Sheet 6

Data Sheet 7

Data Sheet 8

Data Sheet 12

Data Sheet 9

Data Sheet 10

Data Sheet 11

## CREDIT AUTHORSHIP CONTRIBUTION STATEMENT

**Sam Victor:** Writing – original draft, Investigation, Resources, Formal analysis, Methodology, Project administration. **Siddharth Sadasivam:** Conceptualization, Methodology, Data curation, Software, Writing – review and editing. **Gugan Satheesh:** Visualization. **Shravan Aarya Prabhakaran:** Visualization, Validation. **Yuvaraj S:** Supervision, Writing – review and editing. **Subha R:** Supervision.

## ETHICS STATEMENT

This study is a secondary computational analysis of previously published, publicly archived and de-identified 16S rRNA sequence data. No new human samples were collected, no participants were recruited, and no identifiable information was accessed. The data were generated by Jö rissen et al. (2021) and are deposited in the European Nucleotide Archive under PRJEB33591; in that study, ethical approval was obtained from the ethics committee of Antwerp University Hospital, the protocol was registered at ClinicalTrials.gov under NCT03109496 (12 April 2017), and written informed consent was obtained from a parent or legal guardian before sampling. Ethical review and approval were not required for the present analysis in accordance with local legislation and institutional requirements.

## DATA AVAILABILITY STATEMENT

Sequence data are publicly available from the European Nucleotide Archive under study accession PRJEB33591 (Jörissen et al., 2021) and the individual run accessions of Table 1. All benchmarking code, the measured data, the classifier training script, the resource predictor, the analysis and figure-generation scripts, and a `conda` export of the measurement environment are at https://github.com/samvictordr/16S-Amplicon-rRNA-pipeline-protocol. Every repetition is deposited, not only the means reported here, one CSV per arm: the two-sample depth contrast, the within-library depth series, the nine samples measured individually, the ASV sweep, the extended query sweep to 60 000, the strong- and weak-scaling ladders, the nested decomposition and its import isolation, the two cold-cache series and the session comparison, the concurrent-throughput arm, the 280 measurements of the randomised replication, the API-against-CLI comparison, the plugin-set comparison with its taxonomy hashes and memory, the NUMA binding test, the sampling profile with its per-phase timings, the determinism content hashes, and the cohort’s taxonomic composition and diversity. Table S1 names the file each reported quantity derives from. Each classification row records the query count in its `reads` column, which is the column-name collision listed in Table S2.

## ACKNOWLEDGMENTS

This work builds on the QIIME 2, DADA2, SILVA, RESCRIPt and SLURM open-source projects.

