## Supplementary PDF for "Naïve Bayes taxonomic classification cost in QIIME 2 is set by the invocation rather than by sequencing depth or community richness at single-sample scale: a measured resource protocol for 16S rRNA pipelines"

### Supplementary Note S1: the between-session effect

Section 3.9 of the manuscript reports that the same single-sequence classification measured 36.05 s in one session and 32.98 s to 33.47 s in later sessions on the same idle machine, about 7% apart against a within-session standard deviation of 0.11 s. Six candidate causes were tested. Five are excluded and one survives; the full account follows.

We tested six candidate causes. Five are excluded and one survives. The harness wrapper is not responsible (33.33 s under `/usr/bin/time -v` against 33.43 s bare); neither is processor frequency state, since a classification run immediately after saturating all 40 cores for 150 s took 33.12 s against 33.24 s on an idle machine; nor is eviction of the reference by intervening work, since a classification following a 620 s denoising run took 33.11 s. Thread pinning, query artifact and command were identical throughout, and the effect is specific to classification: denoising measured in the fast session reproduced its replication value to 0.55% (619.69 s against 623.12 s). Nor did the software change between the sessions: a profiler was installed into the environment in the interval, and an audit of the environment tree shows it added only its own executable and metadata, upgrading no package and regenerating no bytecode.

The fifth candidate was NUMA placement, and it is the first that survives. This machine has two NUMA nodes and no binding policy was applied, so the scheduler is free to place a process on one node while the pages it reads sit on the other. Forcing that configuration is expensive: binding the process to node 0 while binding its memory to node 1 raises classification from 33.46 s unbound to 38.28 s, 4.82 s or 14.4%, where binding both to the same node costs nothing (33.64 s on node 0, 33.15 s on node 1). The effect is larger than the 2.75 s we are trying to explain, so a partially remote placement would suffice; it can persist for a whole session, because cached library pages are placed once when first read and stay there; and it would fall on classification rather than denoising, since classification's dominant cost is reading many shared-object pages while denoising works mostly in memory it allocates itself. We cannot establish retrospectively where the replication's pages and processes sat, so we do not claim this was the cause. It is, however, the one mechanism consistent with the size of the effect, its persistence and its specificity, and it is the one we would control first. The manuscript's measurement-controls section accordingly recommends binding, which we did not do.

The sixth candidate was page-cache state of the import path, which classification pays and denoising does not. It is the most natural explanation and it is wrong. A cold cache does reproduce the slow session's value almost exactly, 35.83 s against 36.05 s, but it cannot reproduce its persistence: recovery is complete after a single run (35.56 s, then 33.66 s for the next five), whereas the replication measured this condition ten consecutive times at 36.05 s with a standard deviation of 0.11 s. The decisive observation is that each session has its own warm baseline and the cold penalty sits on top of whichever one is in force. In the slow session the cold-cache arm measured 36.08 s warm and 37.99 s cold; in the fast session the same contrast is 33.30 s against 35.83 s. Had the slow session merely been running cold, dropping its cache could not have cost a further 1.90 s. What differs between sessions is the warm state itself.

Of the six candidates, then, five are excluded and NUMA placement is the one consistent with the size of the effect, its persistence across a session and its confinement to classification. We cannot confirm it retrospectively and do not assert it. What we can say is that the effect is real, that it is several times the within-session scatter, and that binding CPU and memory to one node is the control we would apply first and did not.

Expected values for every quantity measured in this work, with the deposited file each is derived from. The *Transfers* column states whether we expect a reader reproducing the protocol on different hardware to recover the same value. Absolute times, the fitted $T_1$, the fixed classification cost and the memory intercept are properties of this node, its filesystem and the reference artifact used, and are not expected to transfer. The ratios are the claims.

Table S1: Expected values for every quantity measured in this work, with the deposited file each is derived from.

| Quantity | This system | Transfers | Source |
|---|---|---|---|
| *Depth contrast, two samples at four cores* | | | |
| Denoise wall, 66 993 reads | 48.84 s | no | `depth_pair.csv` |
| Denoise wall, 1 804 054 reads | 626.14 s | no | `depth_pair.csv` |
| Classify wall, both of the above | 41.02 s / 40.64 s | no | `depth_pair.csv` |
| **Denoise response to 27× depth** | 12.8× | **yes** | `depth_pair.csv` |
| **Classify response, same contrast** | 1.01× | **yes** | `depth_pair.csv` |
| *Within-library depth series* | | | |
| Denoise, 66 816 to 1 804 054 reads | 43.67 s to 623.12 s (14.3×) | ratio | `depth_series.csv` |
| Classify, same range | 40.63 s to 41.18 s (1.01×) | ratio | `depth_series.csv` |
| ASVs recovered | 21 → 58 (2.8×) | yes | `depth_series.csv` |
| *Cost against richness, nine samples at four cores* | | | |
| Denoise against read count | exponent 0.75; 0.67 for the eight shallow alone | ratio | `per_sample.csv` |
| Classify against ASV count | $R^2$ = 0.003, 0.46 s over 5 to 70 ASVs | yes | `per_sample.csv` |
| **Classify, 5-ASV vs 70-ASV community** | 40.81 s / 40.79 s (1.000×) | **yes** | `per_sample.csv` |
| *Fixed cost and query count* | | | |
| Classify wall, 1 ASV → 218 ASVs | 36.20 s to 37.27 s | no | `asv_sweep.csv` |
| Fixed-cost fraction at 218 ASVs | 97.0% | broadly | `asv_sweep.csv` |
| Fitted fixed cost, 1 to 60 000 queries | 36.36 s | no | `query_sweep.csv` |
| Fitted per-query cost | 2.497 ms | no | `query_sweep.csv` |
| Dominance crossover $k^*$ | 14 560 queries (14 437 to 14 714 ) | no | `query_sweep.csv` |
| Measured bracket on $k^*$ | between 15 000 and 20 000 queries | no | `query_sweep.csv` |
| *Decomposition of the fixed cost* | | | |
| Decomposition total | 36.23 s | no | `fixed_cost.csv` |
| Instantiating the `PluginManager` | 16.23 s | broadly | `import_isolation.csv` |
| Reading + deserialising the reference | 2.2 s (under 7%) | no | `import_isolation.csv` |
| Interpreter and `import qiime2` | 0.78 s | broadly | `import_isolation.csv` |
| Action work after the pipeline exists | 14.07 s, not decomposed | — | `fixed_cost.csv` |
| CPU share of the fixed cost | 99.9% | **yes** | `fixed_cost.csv` |
| **Reference on `tmpfs` vs NVMe** | 0.00 s, **CI** −0.07 **to** 0.07 | **yes** | `fixed_cost.csv` |
| Cold-cache penalty, paired | 1.90 s, CI 1.83 to 1.97 | broadly | `cold_cache.csv` |
| Cold-cache penalty, second series | 2.53 s (different session) | broadly | `session_cache.csv` |
| of which in the import path | 2.08 s (82%) | broadly | `session_cache.csv` |
| Recovery after one cold run | complete; 35.56 s then 33.66 s | — | `session_cache.csv` |
| Between-session shift, classification | 2.75 s (7%), unexplained | — | `session_cache.csv` |

| Quantity | This system | Transfers | Source |
|---|---|---|---|
| Between-session shift, denoising | 0.55% | — | probe_tie.csv |
| Cross-node NUMA placement penalty | 4.82 s (14.4%) | broadly | numa_placement.csv |
| Same-node binding, either node | no cost vs unbound | broadly | numa_placement.csv |
| *Installed plugin set* | | | |
| Classification, full distribution (27 plugins) | 33.24 s | no | lean_env.csv |
| Classification, minimal environment (5 plugins) | 18.75 s | no | lean_env.csv |
| **Reduction from a minimal environment** | 43.6% | **broadly** | lean_env.csv |
| Peak memory, full / minimal | 3.10 GiB / 2.22 GiB (−28.3%) | no | lean_env_memory.csv |
| **Taxonomy across the two environments** | **byte-identical** | **yes** | lean_env_hashes.tsv |
| *Concurrent throughput, single-core jobs* | | | |
| Throughput at $N = 40$ | 21.13× the single-job rate | broadly | concurrency.csv |
| Per-job wall, $N = 1 \to 40$ | 37.59 s to 68.31 s (1.82×) | no | concurrency.csv |
| Advantage over four cores in four waves | 2.74× | broadly | concurrency.csv |
| Peak aggregate RSS at $N = 40$ | 115.5 GiB (2.89 GiB to 3.09 GiB per job) | no | concurrency_memory.csv |
| Failures at any level | none | — | concurrency.csv |
| *Randomised replication, ten repetitions,* | 280 *measurements* | | |
| Largest shift vs original, denoising | 0.21 s (0.43%) | — | strong_scaling_rep10.csv |
| Largest shift vs original, ASV sweep | 0.43 s (1.04%) | — | asv_sweep_rep10.csv |
| Mean within-cell SD, original / replicate | 0.19 s / 0.22 s | — | both files |
| Fixed cost remeasured, one sequence | 36.05 s (SD 0.11) | no | asv_sweep_rep10.csv |
| *Thread scaling* | | | |
| Denoise $T_1$ at 66 993 reads | 56.61 s | no | strong_scaling.csv |
| Denoise peak speedup | 1.184× at 8 threads | broadly | strong_scaling.csv |
| **Classify speedup, every $p > 1$** | **below** 1.0 | **yes** | strong_scaling.csv |
| Classify speedup at 40 jobs | 0.782× at 10.5× the CPU | no | strong_scaling.csv |
| USL fit, denoising | $\alpha = 0.818$, $\beta = 0.000\,916$ | no | fitted_models.json |
| *Determinism and memory* | | | |
| **ASV sequences at 1 / 4 / 40 threads** | **identical**, SHA-256 c759115c59be27c5... | **yes** | determinism.tsv |
| **Taxonomy at 1 / 4 / 40 jobs** | **identical**, SHA-256 11b2812c... | **yes** | taxonomy_determinism.tsv |
| Denoise memory model | 1.006 GiB + 2.254 MiB per 1 000 reads, $R^2 = 0.995$ | no | depth_series.csv |
| Classifier peak memory, any condition | 3.11 GiB | no | asv_sweep.csv |

Failed runs are excluded from every fit, but a stage that aborts still records a plausible wall time, so check for them explicitly before trusting a reproduction:

```
awk -F, 'FNR>1 && $NF!=0 {print FILENAME": "$1", exit="$NF}' benchmarks/*.csv
```

Table S2: Pitfalls encountered in applying the protocol, with the symptom that reveals each and the remedy. Step numbers refer to the manuscript's Protocol table.

| **Pitfall** | **Symptom** | **Remedy** |
|---|---|---|
| Unpinned linear-algebra threads | CPU time / wall time > 1.0 at one thread | Pin BLAS pools; verify the ratio is ≈ 1.0 first |
| Logical rather than physical cores | Efficiency reported against a core count the machine lacks | Derive the ladder from `lscpu`, not `nproc` |
| Mismatched classifier region | None at runtime; classification succeeds and returns confidence scores | Read primers from artifact provenance at the step 2 pause point |
| Reads retaining PCR primers | Median merged ASV length ≈ 292 bp rather than 253 bp for V4 | Step 3 pause point; trim with `--p-trim-left-f/r` or exclude the dataset |
| Failed runs recorded as measurements | A rejected classifier still records a time (here 32.45 s) | Record exit status per run; discard non-zero rows before fitting |
| Platform version drift | Pipeline fails outright or will not install (four interface changes including a distribution rename, 2025.7 to 2026.7) | Probe for version-dependent options; record the exact environment and the distribution name |
| Samples differing in more than depth | Composition varies alongside depth | Subsample a single library, as in the within-sample series |
| Unbound NUMA placement | Timings differ between sessions with no change in the command: 7% observed here, and 14% when a cross-node placement is forced deliberately | Bind CPU and memory to one node (`numactl --cpunodebind=N --membind=N`) and report it |
| Reusing one column for two quantities | A classification row's `reads` column holds a query count, not a read count | Keep the instrumentation schema fixed but document the overload; read the stage tag before the column |

Table S3: Experimental arms. Most were run in triplicate; the decomposition, the randomised replication and the plugin-set comparison used ten repetitions and the cold-cache arms five. "Fixed" names the quantity held constant so that the varied quantity is the only source of difference.

| Arm | Varies | Fixed, and why it is informative |
|---|---|---|
| Depth comparison | ERR3444605 vs ERR3444628, 26.9× in depth | Four cores; different participants, so complemented by the within-sample series |
| Within-sample depth series | One library subsampled to six depths by deterministic prefix | Four cores and composition exactly fixed; depth is the only variable |
| Per-sample cost | Each of the nine samples individually | Four cores; cost against each sample's own measured richness |
| ASV sweep | Nested subsets of the pooled union at 1, 10, 25, 50, 100, 218 | One and eight jobs; $k = 1$ isolates the fixed cost |
| Extended query sweep | Nested subsets of 1 to 60 000 read-derived queries | Length fixed at 250 bp, so query count is the only variable |
| Strong scaling | $p \in \{1, 2, 4, 8, 16, 20, 32, 40\}$ cores | Input fixed; $p = 20$ samples the socket boundary |
| Weak scaling | Read count scaled with core count | 1 674 reads per core held constant |
| Output determinism | Denoising at 1, 4 and 40 threads | Representative sequences compared by content hash |
| Fixed-cost decomposition | Each layer of the invocation, timed separately | Ten repetitions in randomised order; classifier on NVMe and on `tmpfs` |
| Cold cache | Page cache dropped immediately before each run | Five cold runs, each paired with the warm run that followed it |
| Concurrent throughput | $N \in \{1, 2, 4, 8, 20, 40\}$ simultaneous single-core jobs | Batch timed to the last finisher; aggregate RSS sampled each second |
| Randomised replication | Strong scaling and ASV sweep re-measured eight weeks later | Ten repetitions, all cells shuffled into one sequence; tests for drift from fixed ordering |
| Invocation split | `PluginManager`, artifact load and pipeline view timed separately | Three repetitions; separates framework import from the reference |
| Sampling profile | One invocation sampled at 50 Hz | Attribution only; the profiler slows the run it samples |
| Python API against CLI | The same work driven both ways | One session, interleaved; isolates the command-line layer |
| Session and cache state | Warm and cold runs of import alone and of the full stage | Cache dropped between; recovery tracked over six runs |
| Installed plugin set | Full distribution against a minimal environment | Interleaved in one session; taxonomy compared by hash |
| NUMA placement | Unbound, bound to either node, and deliberately cross-node | One session; tests a control we had not applied |

Table S4: Community structure. Reads are the number retained in the feature table after denoising. $H'$ is the Shannon index computed on ASV relative abundances.

| Run | Participant | Site | Reads | ASVs | $H'$ | Dominant genus (%) |
|---|---|---|---|---|---|---|
| ERR3444623 | D-005 | middle ear | 46 381 | 5 | 0.005 | *Haemophilus* (99.9) |
| ERR3444633 | D-006 | middle ear | 40 251 | 11 | 1.133 | *Alloiococcus* (61.6) |
| ERR3444606 | D-003 | ear canal | 34 115 | 38 | 1.110 | *Alloiococcus* (67.9) |
| ERR3444597 | D-002 | nasopharynx | 56 463 | 16 | 1.175 | *Moraxella* (82.6) |
| ERR3444605 | D-003 | nasopharynx | 54 959 | 54 | 1.494 | *Moraxella* (58.7) |
| ERR3444642 | D-008 | nasopharynx | 57 836 | 55 | 1.012 | *Moraxella* (66.9) |
| ERR3444628 | D-006 | nasopharynx | 1 070 469 | 58 | 1.311 | *Moraxella* (46.6) |
| ERR3444680 | D-014 | adenoid | 37 649 | 67 | 2.624 | *Haemophilus* (38.0) |
| ERR3444641 | D-008 | adenoid | 56 021 | 70 | 2.578 | *Haemophilus* (32.3) |

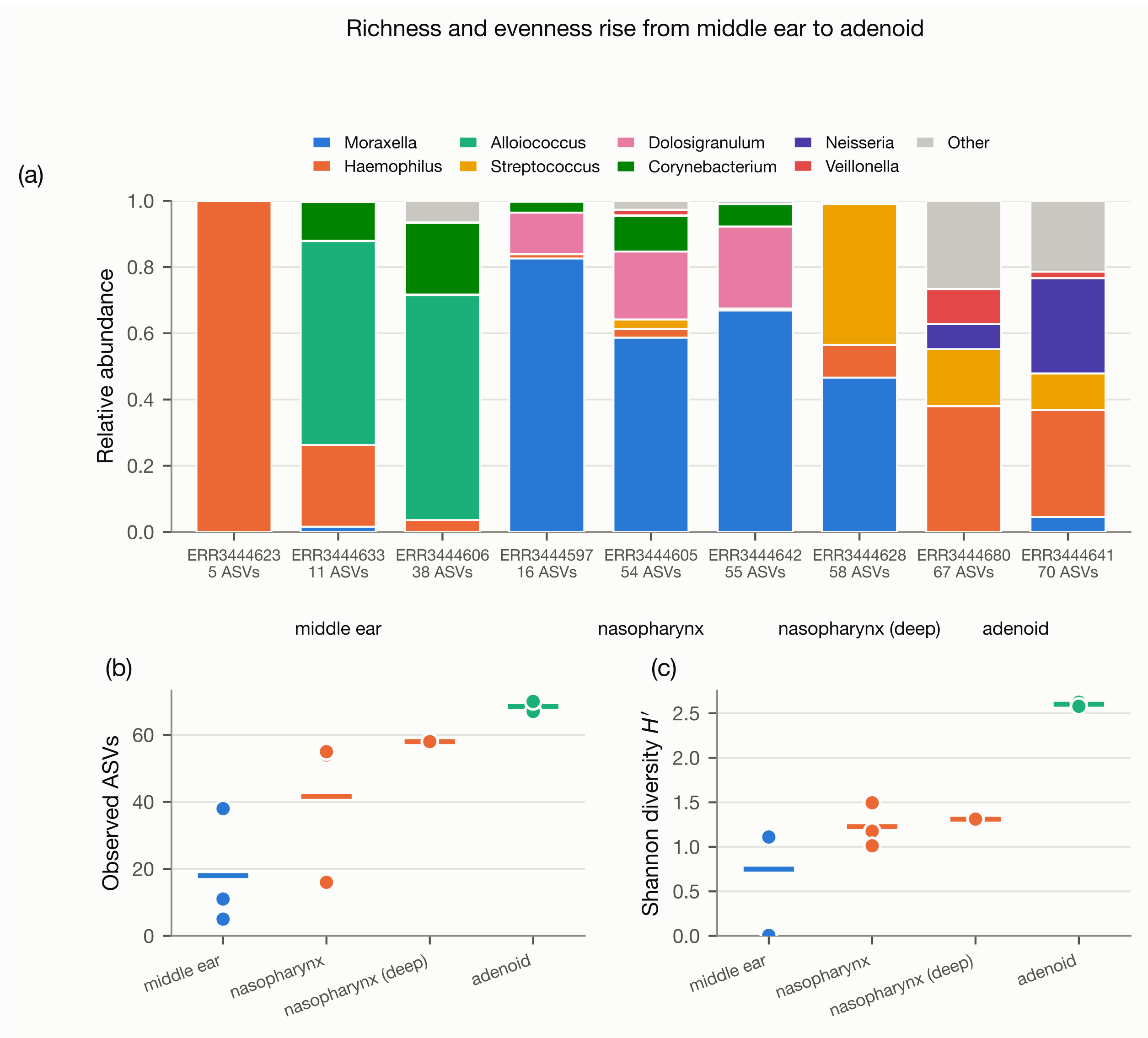


Figure S1: **Community structure across body sites.** **(a)** Genus-level relative abundance per sample, ordered by ASV count and grouped by site; the eight most abundant genera are shown and the remainder pooled as "Other". **(b)** Observed ASVs and **(c)** Shannon diversity by site; bars mark the site mean. The deep sample is shown separately because its sequencing depth is an order of magnitude greater.

Table S5: Depth comparison. Two nasopharyngeal samples from PRJEB33591, at four cores.

| | **ERR3444605** | **ERR3444628** | **Ratio** |
|---|---|---|---|
| Read pairs | 66 993 | 1 804 054 | 26.93 |
| ASVs | 54 | 58 | 1.07 |
| Denoise wall time (s) | 48.84 | 626.14 | 12.82 |
| Denoise peak RSS (GiB) | 1.34 | 5.01 | 3.75 |
| Classify wall time (s) | 41.02 | 40.64 | 0.99 |
| Classify peak RSS (GiB) | 3.11 | 3.11 | 1.00 |

Table S6: Within-sample depth series. One library (ERR3444628) subsampled by deterministic prefix selection; both stages at four cores, means of three repetitions (± SD).

| Read pairs | ASVs | Denoise (s) | Classify (s) |
|---:|---:|---:|---:|
| 66 816 | 21 | 43.67 ± 0.17 | 40.63 ± 0.30 |
| 128 861 | 26 | 59.71 ± 0.25 | 41.24 ± 1.29 |
| 257 722 | 33 | 103.42 ± 6.69 | 40.60 ± 0.31 |
| 451 013 | 35 | 162.72 ± 1.85 | 40.91 ± 0.42 |
| 902 027 | 47 | 308.58 ± 0.21 | 41.00 ± 0.66 |
| 1 804 054 | 58 | 623.12 ± 1.86 | 41.18 ± 0.70 |
| **ratio** | 2.76× | 14.27× | 1.01× |

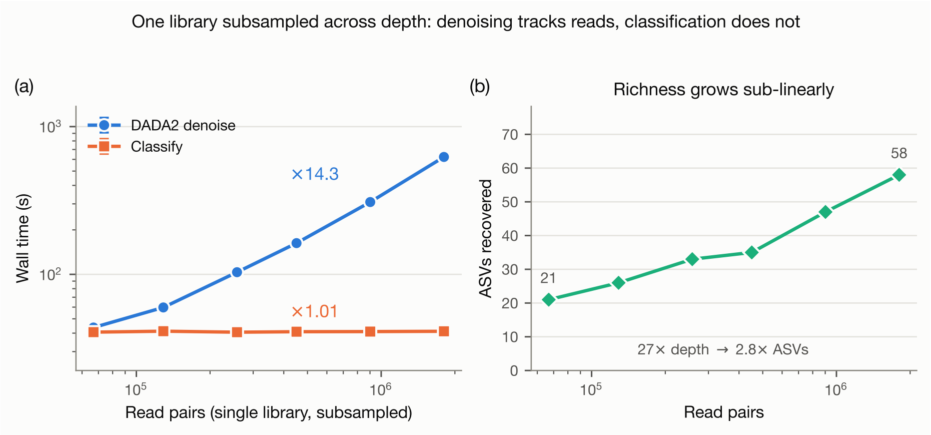


Figure S2: **One library subsampled across a 27-fold depth range. (a)** Wall time for both stages against read count; error bars are ±1 SD over three repetitions, both axes logarithmic. **(b)** ASVs recovered at each depth. Richness grows sub-linearly but substantially, and classification cost does not follow it.

Table S7: Classification wall time against ASV count, means of three repetitions (± SD). Sequences are nested subsets of the 218 distinct representative sequences pooled across the nine samples.

| ASVs | 1 job (s) | 8 jobs (s) |
|---|---|---:|
| 1 | 36.20 ± 0.10 | 38.90 ± 0.12 |
| 10 | 36.07 ± 0.09 | 40.72 ± 0.32 |
| 25 | 36.43 ± 0.09 | 41.26 ± 0.30 |
| 50 | 36.70 ± 0.10 | 42.36 ± 1.03 |
| 100 | 36.95 ± 0.06 | 41.87 ± 0.19 |
| 218 | 37.27 ± 0.08 | 42.05 ± 0.19 |
| *1 job:* intercept 36.25 s, slope 5.21 ms per ASV ($R^2 = 0.86$) | | |

Table S8: Strong scaling. Denoising on ERR3444605 (66 993 read pairs); classification on a 54-sequence ASV set. $S(p)$ is speedup relative to one core, $E(p)$ parallel efficiency.

| | **DADA2 denoise** | | | | **Classify** | | | |
|---|---|---|---|---|---|---|---|---|
| $p$ | Wall (s) | CPU (s) | $S(p)$ | $E(p)$ | Wall (s) | CPU (s) | $S(p)$ | $E(p)$ |
| 1 | 56.61 | 56.5 | 1.000 | 100.0% | 36.68 | 36.6 | 1.000 | 100.0% |
| 2 | 51.23 | 56.7 | 1.105 | 55.3% | 40.94 | 49.1 | 0.896 | 44.8% |
| 4 | 48.90 | 58.4 | 1.158 | 28.9% | 41.03 | 64.3 | 0.894 | 22.3% |
| 8 | 47.81 | 62.1 | 1.184 | 14.8% | 42.57 | 103.9 | 0.862 | 10.8% |
| 16 | 47.88 | 76.3 | 1.182 | 7.4% | 43.11 | 174.7 | 0.851 | 5.3% |
| 20 | 47.85 | 81.8 | 1.183 | 5.9% | 44.13 | 229.3 | 0.831 | 4.2% |
| 32 | 47.97 | 107.7 | 1.180 | 3.7% | 46.87 | 377.5 | 0.783 | 2.4% |
| 40 | 48.50 | 131.5 | 1.167 | 2.9% | 46.92 | 383.7 | 0.782 | 2.0% |

Table S9: Paired comparison of the two query sets at the five query counts they share. The ASV set is the pooled representative sequences of the nine samples, of median length 253 bp. The read set is dereplicated R1 reads from ERR3444628 trimmed to a fixed 250 bp. Values are means of three repetitions. The mean absolute difference is 0.65%, the median 0.56% and the range −0.7 to 1.8%. Differences run in both directions, so the read-derived pool shows no systematic offset from real ASV sets over the range where the two can be compared.

| **Jobs** | **Queries** | **ASV set** (s) | **Read set** (s) | **Difference** |
|---|---|---|---|---|
| 1 | 1 | 36.20 | 36.49 | 0.8% |
| 1 | 10 | 36.07 | 36.47 | 1.1% |
| 1 | 50 | 36.70 | 36.77 | 0.2% |
| 1 | 100 | 36.95 | 37.17 | 0.6% |
| 1 | 218 | 37.27 | 37.24 | −0.1% |
| 8 | 1 | 38.90 | 39.03 | 0.3% |
| 8 | 10 | 40.72 | 40.94 | 0.5% |
| 8 | 50 | 42.36 | 42.05 | −0.7% |
| 8 | 100 | 41.87 | 42.63 | 1.8% |
| 8 | 218 | 42.05 | 42.19 | 0.3% |

Table S10: Predicted against measured, for eight samples held out of the calibration entirely. Negative errors are under-predictions. The classification prediction is the constant 40.93 s, which is why its column has no separate predicted entry.

| **Reads** | **Denoising wall time** | | | **Denoising peak memory** | | | **Classification** | |
|---|---|---|---|---|---|---|---|---|
| | measured | predicted | error | measured | predicted | error | measured | error |
| 44 604 | 40.02 s | 26.97 s | −32.6% | 1.15 GiB | 1.10 GiB | −4.4% | 41.21 s | −0.69% |
| 54 938 | 46.30 s | 31.97 s | −30.9% | 1.22 GiB | 1.13 GiB | −7.6% | 40.75 s | +0.43% |
| 62 444 | 42.25 s | 35.49 s | −16.0% | 1.14 GiB | 1.14 GiB | +0.6% | 40.81 s | +0.30% |
| 62 686 | 58.36 s | 35.61 s | −39.0% | 1.26 GiB | 1.14 GiB | −9.4% | 41.00 s | −0.17% |
| 66 212 | 47.53 s | 37.23 s | −21.7% | 1.26 GiB | 1.15 GiB | −8.5% | 40.90 s | +0.08% |
| 66 993 | 49.05 s | 37.59 s | −23.4% | 1.34 GiB | 1.15 GiB | −13.7% | 41.03 s | −0.24% |
| 70 630 | 54.05 s | 39.25 s | −27.4% | 1.24 GiB | 1.16 GiB | −6.3% | 41.13 s | −0.49% |
| 82 257 | 75.22 s | 44.45 s | −40.9% | 1.36 GiB | 1.19 GiB | −13.0% | 40.79 s | +0.34% |
| mean absolute error | | | 29.0% | | | 7.9% | | |

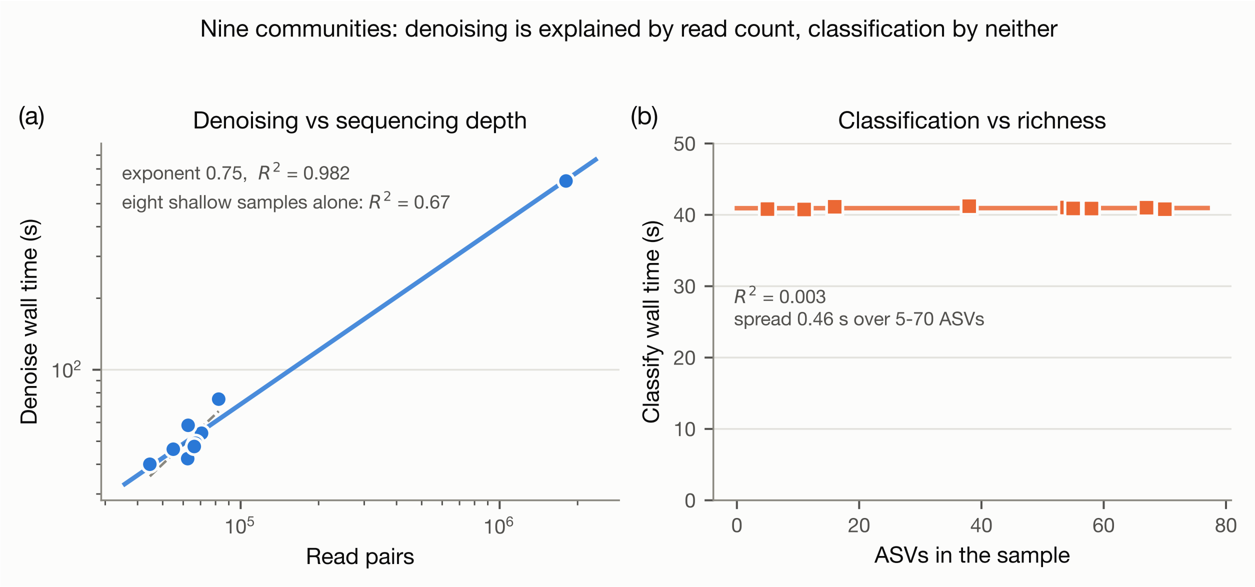


Figure S3: **Nine communities measured individually. (a)** Denoising wall time against read count, logarithmic axes, with the fitted power law. The annotation reports both the pooled fit and the fit restricted to the eight shallow samples, since the pooled value is influenced by the single deep sample. **(b)** Classification wall time against the number of ASVs in the sample.

Table S11: Measured throughput under sample-level parallelism. $N$ identical single-core classifications were started simultaneously and the batch timed to the last finisher; three repetitions per level. Throughput is $N$ divided by batch wall time.

| $N$ | **Batch wall** (s) | **Throughput** ($s^{-1}$) | **Per-job mean** (s) | **Relative throughput** |
|---|---|---|---|---|
| 1 | 37.60 | 0.027 | 37.59 | 1.00× |
| 2 | 40.09 | 0.050 | 39.50 | 1.88× |
| 4 | 43.16 | 0.093 | 42.16 | 3.50× |
| 8 | 45.46 | 0.176 | 43.52 | 6.63× |
| 20 | 53.58 | 0.373 | 51.31 | 14.07× |
| 40 | 71.40 | 0.561 | 68.31 | 21.13× |

Table S12: Choosing the unit of classification. Query count means *distinct* representative sequences presented in one call. Thresholds are those measured on this node with the 62 MiB reference and should be re-derived elsewhere; the manuscript's protocol is what re-derives them. The memory figures are the classification stage's measured peak rounded up for a modest margin, and apply to a job doing classification alone; the manuscript's worked example asks for more because it covers both stages in one job and doubles for margin.

| **Invocation unit** | **Typical query count** | **Allocation** | **Why** |
|---|---|---|---|
| Per sample | 5–100 | One job, 4 GiB (3 GiB in a minimal environment), budget the fixed cost once per sample | Entirely fixed-cost; threads only add overhead. Cost × sample count, so batching saves most here |
| Per batch or cohort union | 100 to 5 000 | One job, 4 GiB (3 GiB in a minimal environment), budget one fixed cost for the whole batch | Union grows sub-linearly in sample count; the fixed cost is paid once instead of per sample |
| Large cohort union | > 5 000 | Multiple jobs; re-measure before committing | Parallelism begins to pay above ≈ 5 000, and above $k^* \approx 14\,560$ the per-query term dominates the fixed cost |

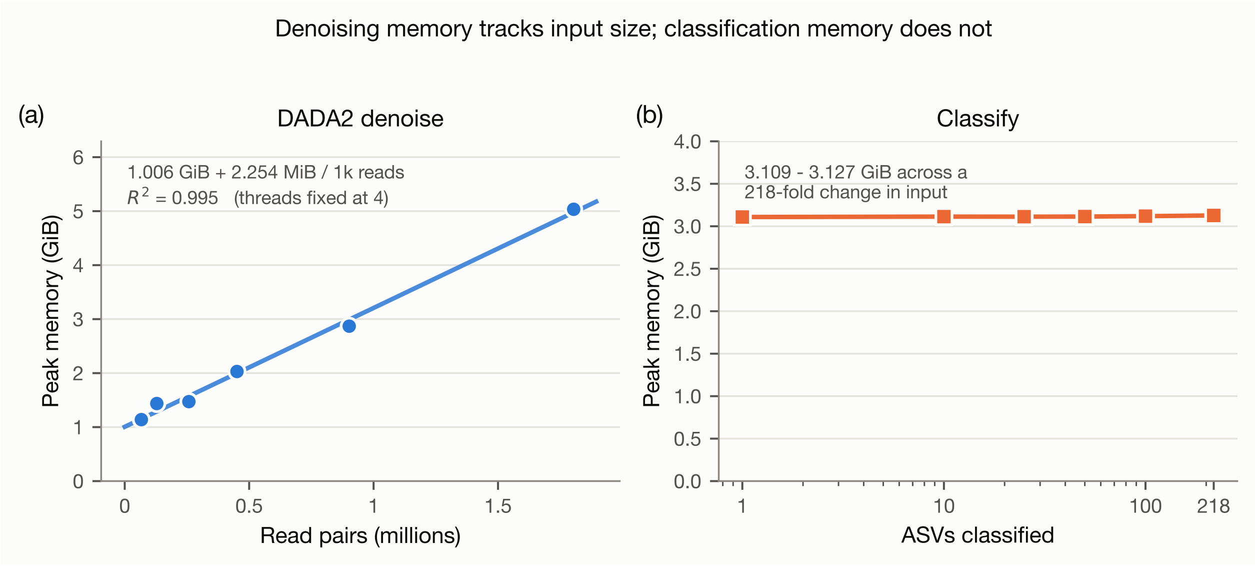


Figure S4: **Denoising memory follows input size; classification memory does not. (a)** Denoising peak memory against read count over the within-sample depth series, in which thread count is held fixed at four, with the fitted memory model of the manuscript. The weak-scaling series is not used here for the reason given in the manuscript's Memory section. **(b)** Classification peak memory against ASV count, which varies by 18 MiB across a 218-fold change in input. Both panels have zero-based vertical axes.

### Description of Data Sheets 1 to 12

Data Sheets 1 to 12 are the measurement records behind the figures and tables of this document and of the manuscript. Each is a plain-text table with a header row. The file names in the second column are the names the same files carry in the project repository, where they sit alongside the scripts that produced them.

Seven of the sheets (2, 4, 5, 8, 9, 11 and 12) share one instrumentation schema, written by the measurement harness once per run: `stage`, a tag naming the pipeline stage and the arm; `threads`, the thread or job count; `reads`, the input size; `wall_sec`, `user_sec` and `sys_sec`, elapsed and CPU time in seconds; `peak_rss_kb`, peak resident set size in kibibytes; `ctx_voluntary` and `ctx_involuntary`, context switch counts; `fs_inputs` and `fs_outputs`, filesystem block counts; `timestamp`, the UTC start time; and `exit_code`, the process exit status.

Two cautions apply when reading these seven. First, the `reads` column is overloaded: on a `denoise` row it holds a read-pair count, but on a `classify` row it holds the number of query sequences, so the `stage` tag must be read before the column. Second, rows with a non-zero `exit_code` record a run that aborted, and such a run still carries a plausible wall time; these rows are excluded from every fit reported here and should be excluded from any reproduction, as noted at the foot of Table S1.

| Data Sheet | Repository file | Contents |
|---|---|---|
| 1 | `alpha_diversity` | Per-sample richness and evenness for the nine libraries: retained reads, observed ASVs, Shannon index, Simpson evenness, and the dominant genus with its relative abundance. Nine rows, one per library. The source for Table S4 and for panels (b) and (c) of Figure S1 |
| 2 | `asv_sweep` | ASV sweep. Classification of nested subsets of the pooled ASV union at 1, 10, 25, 50, 100 and 218 sequences, at one and eight jobs, in triplicate. Thirty-six rows. The source for Table S7, and the arm from which the fixed cost is read at $k = 1$ |
| 3 | `composition` | Taxonomic composition in long format: one row per sample, taxonomic level and taxon, with read count and relative abundance, at phylum and genus level. 282 rows. The source for panel (a) of Figure S1 |
| 4 | `depth_pair` | Depth comparison. Denoising and classification of ERR3444605 against ERR3444628, which differ 26.9-fold in sequencing depth, at four cores. Twelve rows. The source for Table S5 |
| 5 | `depth_series` | Within-sample depth series. One library (ERR3444628) subsampled to six depths by deterministic prefix, at four cores, holding composition exactly fixed so that depth is the only variable. Thirty-six rows. The source for Table S6, Figure S2 and the denoising memory model of panel (a) of Figure S4 |
| 6 | `determinism` | Output determinism for denoising. SHA-256 digest of the representative sequences produced at 1, 4 and 40 threads, with the ASV count. Three rows |

| Data Sheet | Repository file | Contents |
|---|---|---|
| 7 | `fitted_models` | Fitted model parameters, one row per stage, parameter and value: single-thread times, input-size exponents, Amdahl serial fractions, Universal Scalability Law coefficients $\alpha$ and $\beta$, and the corresponding $R^2$ values and validity flags. 180 rows. Flattened from the JSON file of the same name in the repository |
| 8 | `per_sample` | Per-sample cost. Denoising and classification of each of the nine libraries individually at four cores, in triplicate, giving cost against each sample's own measured richness. Fifty-four rows. The source for Figure S3 |
| 9 | `strong_scaling` | Strong scaling. Both stages at 1, 2, 4, 8, 16, 20, 32 and 40 cores with the input held fixed; $p = 20$ samples the socket boundary. Forty-eight rows. The source for Table S8 |
| 10 | `taxonomy_determinism` | Output determinism for classification. SHA-256 digest of the taxonomy assignments produced at 1, 4 and 40 jobs, with the assignment count. Three rows |
| 11 | `weak_scaling` | Weak scaling. Read count scaled with core count at 1 674 read pairs per core, so that per-core work is held constant. Twenty-four rows |
| 12 | `query_sweep` | Extended query sweep. Classification of nested subsets of 1 to 60 000 read-derived queries, each trimmed to a fixed 250 bp so that query count is the only variable, at one and eight jobs, in triplicate. Ninety rows. The source for the crossover analysis and for the per-query slope reported in the manuscript |
